# Hairpin Architecture and Extended Sequence Context Shape APOBEC3 Mutagenesis in Monkeypox Virus at Different Genomic Scales

**DOI:** 10.64898/2026.10.03.756454

**Authors:** Alexandra V. Madorskaya, Fedor M. Kazanov, Danila A. Bogdanov, Gennady V. Ponomarev, Anna E. Smolnikova, Daria V. Garshina, Evgenii V. Matveev, Ruslan Kh. Abasov, Cem Azgari, Ogun Adebali, Dmitry N. Ivankov, Mikhail S. Gelfand, Marat D. Kazanov

## Abstract

The 2022 global mpox outbreak was characterized by an unusual accumulation of more than fifty APOBEC-signature substitutions in MPXV genomes. A previous study suggested that these substitutions preferentially occur within and near hairpins or cruciforms – stem-loop structures formed by inverted repeats. Here, we show that this conclusion resulted from an inappropriate statistical analysis and, using two substantially larger datasets, demonstrate that APOBEC-signature mutations are specifically enriched at TC motifs located at the 3′ ends of hairpin loops, but not in other regions of hairpins/cruciforms or in their vicinity. We further identified a preference for stem-loop structures with longer stems and shorter loops, consistent with patterns previously reported for the APOBEC mutagenesis in cancer genomes. Our analysis also revealed a positive association between APOBEC-signature mutation propensity and hairpin stability in single-stranded DNA. Nevertheless, stem-loop structures with long stems and short loops are relatively rare in the MPXV genome, limiting their contribution to the genome-wide distribution of APOBEC-signature mutations to a predominantly local effect. In contrast, extended nucleotide context downstream of the TC motif showed strong genome-wide predictive value, with a marked preference for guanine-rich trinucleotide immediately downstream. Analysis of the upstream nucleotide context was consistent with the mutational specificity of APOBEC3A. Together, these results identify distinct structural and sequence determinants of the APOBEC-associated mutagenesis in MPXV and show that local hairpin/cruciform architecture and extended nucleotide context contribute at different genomic scales.

## Introduction

Monkeypox virus (MPXV) is a double-stranded DNA virus belonging to the Poxviridae family that has caused a multicountry outbreak in 2022 [1]. Although orthopoxviruses usually evolve slowly (∼1–2 substitutions per genome per year), 2022 MPXV genomes showed an unusual excess of more than 50 substitutions dominated by C→T/G→A changes in TC/ GA contexts [2]. This pattern is consistent with the activity of APOBEC3 cytidine deaminases, intrinsic antiviral restriction factors that deaminate cytosines in single-stranded nucleic acids [3]. The excess of APO-BEC-signature mutations points to the host-driven editing during sustained human transmission, although the extent and substrate of the APOBEC activity in MPXV remain to be clarified. Recently, Dobrovolna *et al*. [4] reported that APOBEC-signature mutations observed during the 2022 monkeypox virus outbreak are enriched in predicted cruciform structures of the viral genome formed by inverted repeats (IRs) generating two opposite hairpins. While the preference of APOBEC mutations for specific secondary structures formed in DNA has been reported before [5], we found that the statistical support for this conclusion is limited and merits further examination.

Indeed, the authors based this conclusion on the observation that “*63.9% of the SNPs were found to arise within IRs*” [4]. However, when replicating the hairpin/cruciform predictions by detecting inverted repeats using the same bioinformatic tool and parameter set described in the study, we found that IRs span 52.4% of the MPXV genome. This implies that even random mutations would be expected to overlap predicted stem-loop structures approximately half of the time. Moreover, the authors’ calculation of 63.9% (39 out of 61 mutations) included seven mutations located merely ‘close’ to, but not within, the IRs, which may substantially bias the statistical interpretation. When applying the binomial test to the enrichment reported in the original study, specifically, 32 out of 61 mutations occurring within IRs, the resulting p-value is 0.55, indicating no statistically significant evidence to reject the null hypothesis of a random distribution of mutations across the MPXV genome.

Other concerns arise from the analysis treating entire hairpins/cruciforms as uniform structures, whereas previous studies have shown that APOBEC-induced mutations are predominantly enriched in hairpin loops, particularly at cytosines located at the 3′ terminus of the loop, both within and outside the TC motif [6]. Moreover, many detected IRs overlap and represent mutually exclusive stem-loop structures, making their simultaneous presence in vivo unlikely and further limiting biological interpretability. Taking this into account, we rigorously reevaluated the Dobrovolna et al.’s hypothesis and found that the number of mutations in the 2022 outbreak dataset provides limited statistical power for assessing structural enrichment. Reassessment using two new, larger datasets showed strong preference of APOBEC-like mutations not to hairpin/cruciform structures or their vicinity as a whole, but specifically to the TC motifs at the 3’ ends of the hairpin loops, consistent with previous studies of the APOBEC activity in cancer [7]. Similar to finding in cancer genomes, the optimal substrate for APOBEC were found stem-loop structures with a longer stem and a short loop. Our findings demonstrate a non-random distribution of candidate APOBEC-induced mutations in the MPXV genome and provide a basis for predicting, to some extent, the genomic sites susceptible to mutation during interactions between the virus and the host immune system following zoonotic transmission.

## Results

### The 2022 MPXV Outbreak Mutation Dataset Shows No Significant Enrichment of APOBEC-Signature Mutations in Hairpin/ Cruciform-Associated Structural Contexts

To address the question of whether APOBEC-signature mutations preferentially occur in hairpin/cruci-form-associated structural contexts within IR regions of the MPXV genome, we performed an independent analysis using the 2022 outbreak mutation dataset analyzed by Dobrovolna et al. [4]. In addition to analyzing the unfiltered set of overlapping IRs, we implemented four alternative strategies (Fig. 1a) to resolve IR overlaps and identify biologically plausible subsets of stem-loop structures: (1) retaining only the most stable structure within each group of overlapping IRs; (2) retaining non-overlapping structures with the highest stability scores; (3) retaining non-overlapping structures that collectively maximize the total length (representing the upper bound of the genomic coverage); and (4) retaining the single shortest structure per overlapping group (representing the lower bound of the coverage). These approaches allowed us to examine both biologically plausible configurations (strategies 1 and 2) and extreme scenarios of genomic coverage (strategies 3 and 4).

**Figure 1.**
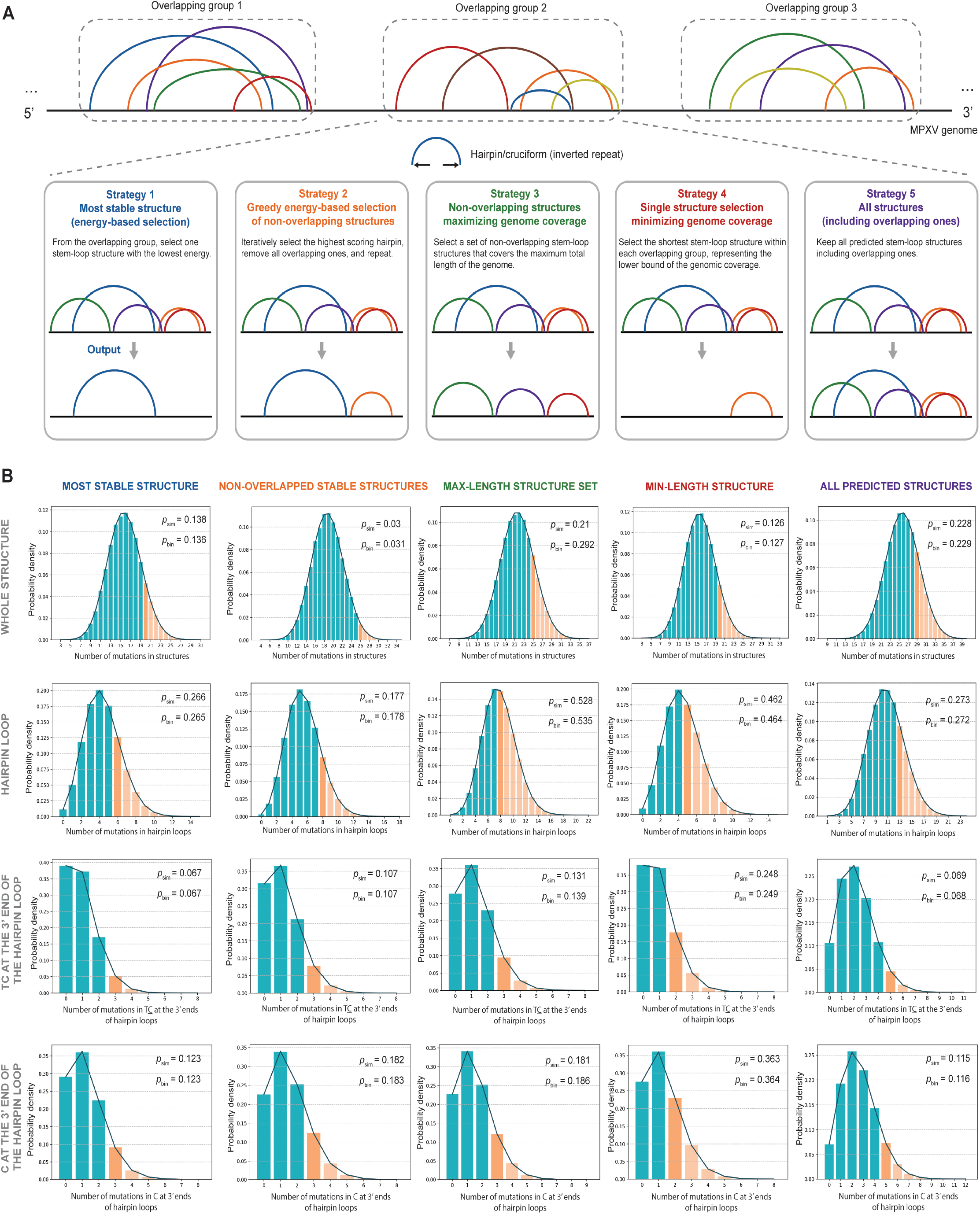
APOBEC3 mutation enrichment analysis in stem-loop structures using the 2022 MPXV outbreak mutation dataset. (a) Strategies for resolving overlapping inverted repeats (IRs) and defining subsets of stem-loop structures. (b) Enrichment analysis of APOBEC3-signature mutations across different structural contexts, including entire stem-loop structures, hairpin loops, cytosines within TC motifs at the 3′ ends of loops (canonical APOBEC hotspots), and cytosines at the 3′ ends of loops irrespective of TC context (extended APOBEC signature). Orange bars indicate observed mutation counts. The *p*-values obtained from Monte Carlo simulations and the binomial test are denoted by *p*_*sim*_ and *p*_*bin*_, respectively.

We quantified the mutation enrichment using two complementary approaches: (i) Monte Carlo simulations, in which mutation positions were randomized across TC/GA motifs in the MPXV genome (preserving the mutation count) and enrichment was recalculated over 100,000 iterations to generate empirical *p*-values; and (ii) binomial testing, comparing the observed number of mutations within predicted stem-loop structures to the expected number based on the genomic coverage. Analyses were performed separately for (a) entire stem-loop structures, (b) hairpin loops, (c) cytosines within TC motifs located at the 3′ ends of hairpin loops (the canonical APOBEC hotspot), and (d) cytosines at the 3’ ends of hairpin loops irrespective of the TC motif (extended APOBEC signature [6]).

Our results are summarized in Fig. 1b. Across all four structure selection strategies, as well as the original unfiltered set of IRs, we observed no statistically significant enrichment of mutations within either entire structures or hairpin loops. Both simulation-based analyses and binomial testing failed to identify statistically significant enrichment. Following the Benja-mini–Hochberg correction for multiple testing, all adjusted p-values exceeded the conventional significance threshold of 0.05, and none of the tested structural contexts were statistically significant. Notably, the strongest trend toward enrichment was observed for mutations in TC motifs at the 3′ ends of hairpin loops. However, the limited number of mutations in this dataset resulted in insufficient statistical power to draw firm conclusions. To address this limitation, we next analyzed a larger collection of West African MPXV genomes [8] and used the ancestral state reconstruction to account for independent mutational events.

### Phylogeny-Informed Analysis Reveals Significant Structural Enrichment of APOBEC-Signature Mutations in West African MPXV Genomes

To increase the statistical power of the analysis, we compiled an additional dataset of MPXV genomes isolated in Nigeria and Cameroon between 2018 and 2023 [8]. This recently published dataset comprises more than one hundred MPXV genomes and shows evidence of APOBEC-mediated mutational activity. We constructed a multiple sequence alignment of the genomes (see Data Availability) and identified positions containing substitutions. The distribution of 3-nucleotide contexts for these substitutions is shown in Fig. 2a. Mutations consistent with the APOBEC mutational signature were predominant, with 5,283 substitutions out of 9,216 total substitutions exhibiting an APOBEC signature. The highest mutation frequency was observed in the TCA triplet, followed by TCG, whereas TCT occurred at approximately half the frequency of the most prevalent triplet, and TCC was the least frequent. This pattern is consistent with previously reported APOBEC trinucleotide context frequencies in cancer studies [9] and further supports the APOBEC-mediated origin of these substitutions.

**Figure 2.**
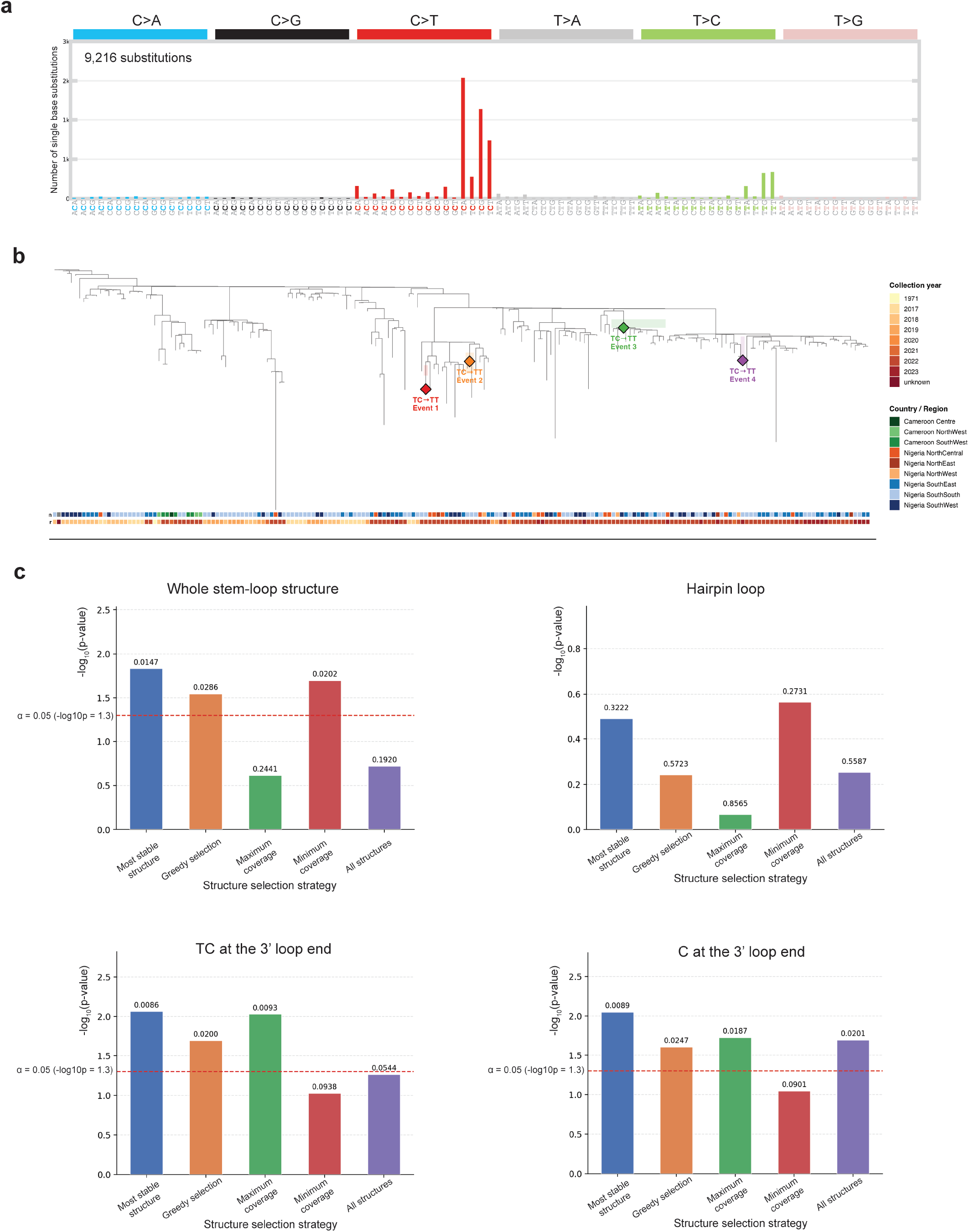
APOBEC-signature mutations in the West African MPXV dataset. (a) Trinucleotide mutation spectrum derived from MPXV genomes collected in Nigeria and Cameroon between 2018 and 2023. The y-axis shows the number of substitution positions for each trinucleotide context. (b) Independent occurrences of the TC→TT substitution at genomic position 35749 (relative to NC_063383.1) inferred by ancestral state reconstruction across the analyzed MPXV genomes. The colored diamond marks these substitutions, which were inferred to have arisen independently in multiple lineages. Branch annotations indicate collection year and geographic origin. (c) Statistical significance of enrichment analyses for APOBEC-signature mutational events in different hairpin/cruciform-associated structural contexts. Bars represent −log10(*p*) values and the dashed red line indicates the significance threshold (*p* = 0.05).

Inspection of the multiple sequence alignment suggested that some substitutions arose independently in different viral lineages (Fig. 2b). Because mutations arising independently at the same genomic position represent independent mutational events, accounting for them can further increase the statistical power of the analysis [10]. To estimate the number of independent events at each position, we reconstructed a phylogenetic tree and inferred ancestral states for all internal nodes. Using the reconstructed evolutionary history, we calculated the number of independently occurring substitutions at each genomic position (see Methods). This analysis identified a total of 1089 independent substitution events for 838 positions with detected APOBEC-like substitutions.

We then repeated the enrichment analysis described in the previous section using independent substitution events inferred by the ancestral reconstruction of the West African dataset. In contrast to the analysis of the 2022 outbreak dataset, APOBEC-signature mutations showed statistically significant enrichment in specific hairpin/cruciform-associated structural contexts (Fig. 2c, Fig. S1). The strongest signal was observed for the TC motif located at the 3’ ends of hairpin loops. Significant enrichment was also observed for cytosines located at the 3’ ends of hairpin loops. However, because this feature is not independent of the TC motif, the enrichment likely reflects the same underlying signal (this was analyzed separately, see below). Weaker enrichment signals were detected for hairpin loops and stem-loop structures considered as a whole. To determine which structural features contribute independently to these enrichment signals, we decomposed stem-loop structures into individual structural components and analyzed each component separately, as described in the following subsections.

### APOBEC-like Mutation Enrichment in Hairpin/Cruciform-Associated Structural Contexts is Robust to Variation in IR Detection Parameters

To assess the robustness of our findings, we evaluated their sensitivity to variations in the parameters used to detect inverted repeats. As a preliminary analysis, we examined how the number of predicted structures and the percentage of genome coverage by IRs vary under different constraint settings, including the minimum and maximum stem length, loop length, and the number of mismatches. First, we estimated how the number of predicted structures and their genome coverage increased as the maximum stem length threshold is relaxed. We recalculated the set of inverted repeats while increasing the maximum stem length threshold from 6 bp, the default minimum stem length, to 50 bp. Fig. 3a shows that once the maximum allowed stem length reaches 9 bp, further increases in the maximum allowed stem length result in only a negligible increase in the number of predicted structures, suggesting that exploring values beyond the default setting of 30 bp is unnecessary and that a lower default may be more appropriate.

**Figure 3.**
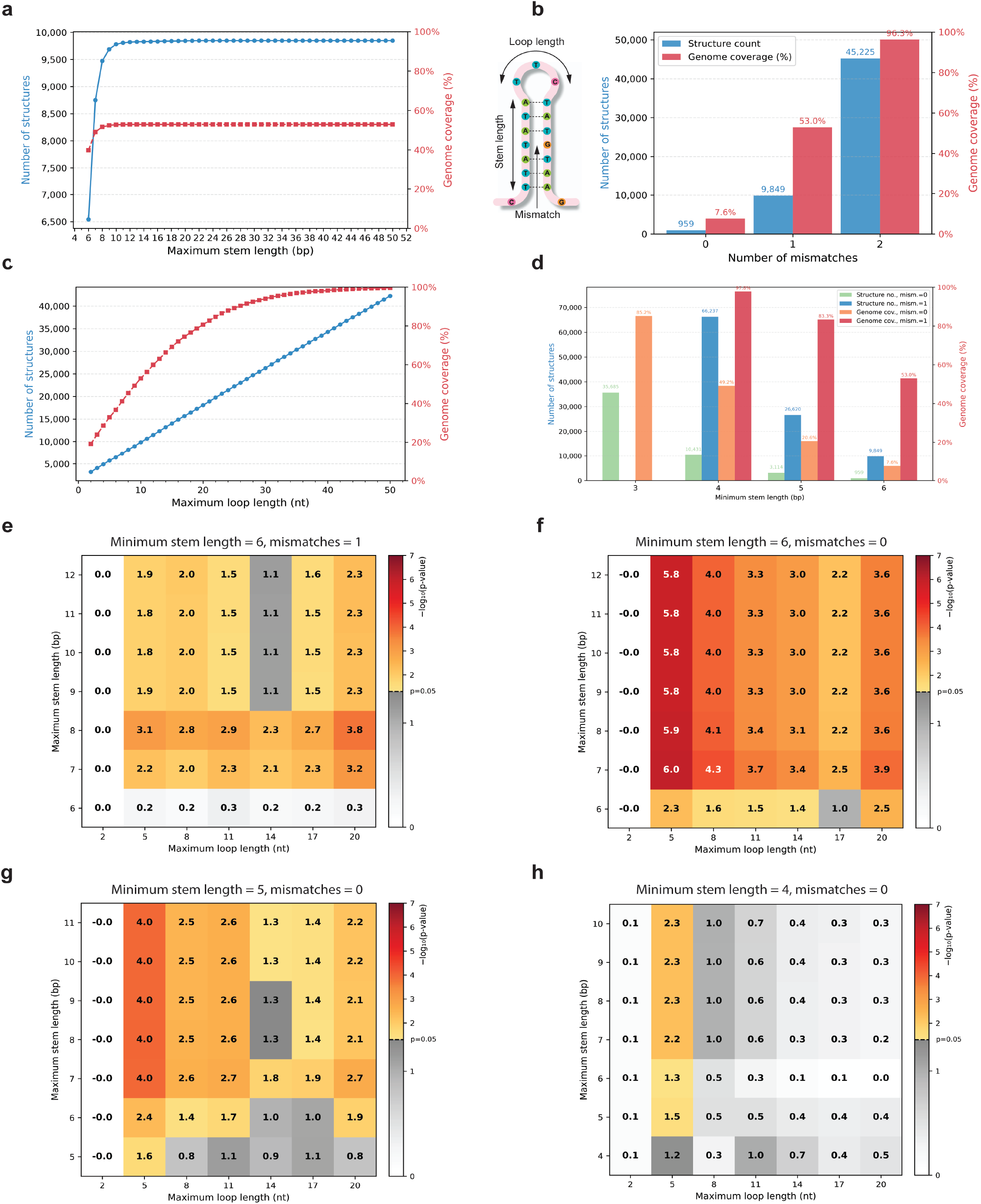
Sensitivity of APOBEC-signature mutation enrichment to IR detection parameters. Number of predicted stem-loop structures in the MPXV genome and percentage of genome coverage by structures as a function of (a) the maximum allowed stem length, (b) the number of allowed mismatches in the stem, (c) the maximum allowed loop length, and (d) the minimum stem length threshold with zero or one allowed mismatch. (e–h) Statistical significance of enrichment analyses for APOBEC-signature mutational events occurring at TC motifs located at the 3′ ends of hairpin loops across different parameter settings for inverted repeat prediction. For groups of overlapping stem-loop structures, analyses were performed using the most stable structure resolution strategy. Heatmaps show −log10(*p*) values for combinations of maximum stem length and maximum loop length. Analyses were performed with minimum stem lengths of 6 nt and 1 mismatch (e), 6 nt and 0 mismatches (f), 5 nt and 0 mismatches (g), and 4 nt and 0 mismatches (h). Gray shading indicates non-significant values (*p* ≥ 0.05).

We further calculated the number of predicted structures and their genome coverage for none, one, and two mismatches within the stem (Fig. 3b). When two mismatches are allowed, both the number of predicted structures and their genome coverage – 45,225 and 96.3%, respectively – become unreasonably high, reducing the ability to detect differences in mutation enrichment between IR and non-IR regions. Thus, we subsequently restricted the maximum number of mismatches to one, resulting in a more reasonable number of predicted structures (9,849) and the genome coverage of 53%. Next, we examined variation in the loop length and found that when the loop length exceeds 20 nt, the genome coverage reaches saturation (>80%), and the number of predicted structures surpasses 20,000 (Fig. 3c). Accordingly, the maximum loop length was limited to 20 nt for subsequent calculations.

Next, we evaluated reasonable values for the minimum stem length. For short stems, the number of allowed mismatches strongly influences the number of predicted structures. We therefore assessed suitable minimum stem length constraints for both none and one mismatch. As shown in Fig. 3d, with one allowed mismatch, a minimum stem length of 6 nt represents a reasonable threshold, as lower values result in predicted structures covering over 80% of the genome. In the absence of mismatches, the threshold can be reduced to 4 nt, corresponding to 49.2% genome coverage, while further reduction to 3 nt again increases genome coverage beyond 80%.

Using the defined parameter ranges, we evaluated, as described above, the enrichment of APOBEC-like mutations in the considered structural features across all parameter values (Figs. S2-S6). Stem-loop structures were predicted using the EMBOSS package [11], as Palindrome Analyzer [12], used in the original study by Dobrovolna et al. [4], could not identify structures across the required range of parameter values. The propensity for TC motifs located at the 3′ end of the hairpin loop (Figs. 3e–h) showed the most significant p-values and remained robust across variations in the IR prediction parameters. It can be seen that statistically significant estimates become stronger under more stringent constraints – i.e., the most significant values are observed when mismatches are not allowed and the minimum stem length is increased, supporting the validity of the observed effect.

### APOBEC-like Mutation Enrichment in the MPXV Genome Is Associated with Specific Structural Geometries and Predominantly Localized to TC Motifs at the 3′ Ends of the Loops

We next investigated whether the observed enrichment depends on the secondary structure geometry, including stem length and loop length, as well as on the position of the target motif within the stem-loop structure. To focus on the parameter range showing the strongest enrichment signal, we restricted our analysis to stem-loop structures without stem mismatches and limited the maximum stem length to 12 bp, as Fig. 3a indicates that increasing the maximum stem length beyond 9-10 bp has little effect on the number of predicted structures. Predicted structures were then stratified according to the stem and loop length, and enrichment of APOBEC-signature substitutions was evaluated for each structural category. As in the previous analyses, enrichment was evaluated using multiple strategies for resolving overlapping IRs.

The strongest enrichment of APOBEC-signature substitutions was observed within a relatively narrow and clustered range of hairpin geometries characterized by long stems and short loops (Fig. 4a, Fig. S7). Specifically, statistically significant enrichment of APO-BEC-like substitutions at TC motifs and cytosines located at the 3′ ends of hairpin loops was detected in stem-loop structures with 3 nt loops and 5-7 bp stems, and in structures with 4 nt loops and 5-6 bp stems. Enrichment was more pronounced for 3 nt loops than for 4 nt loops, with the strongest signal observed for structures with 3 nt loops and 7 bp stems.

**Figure 4.**
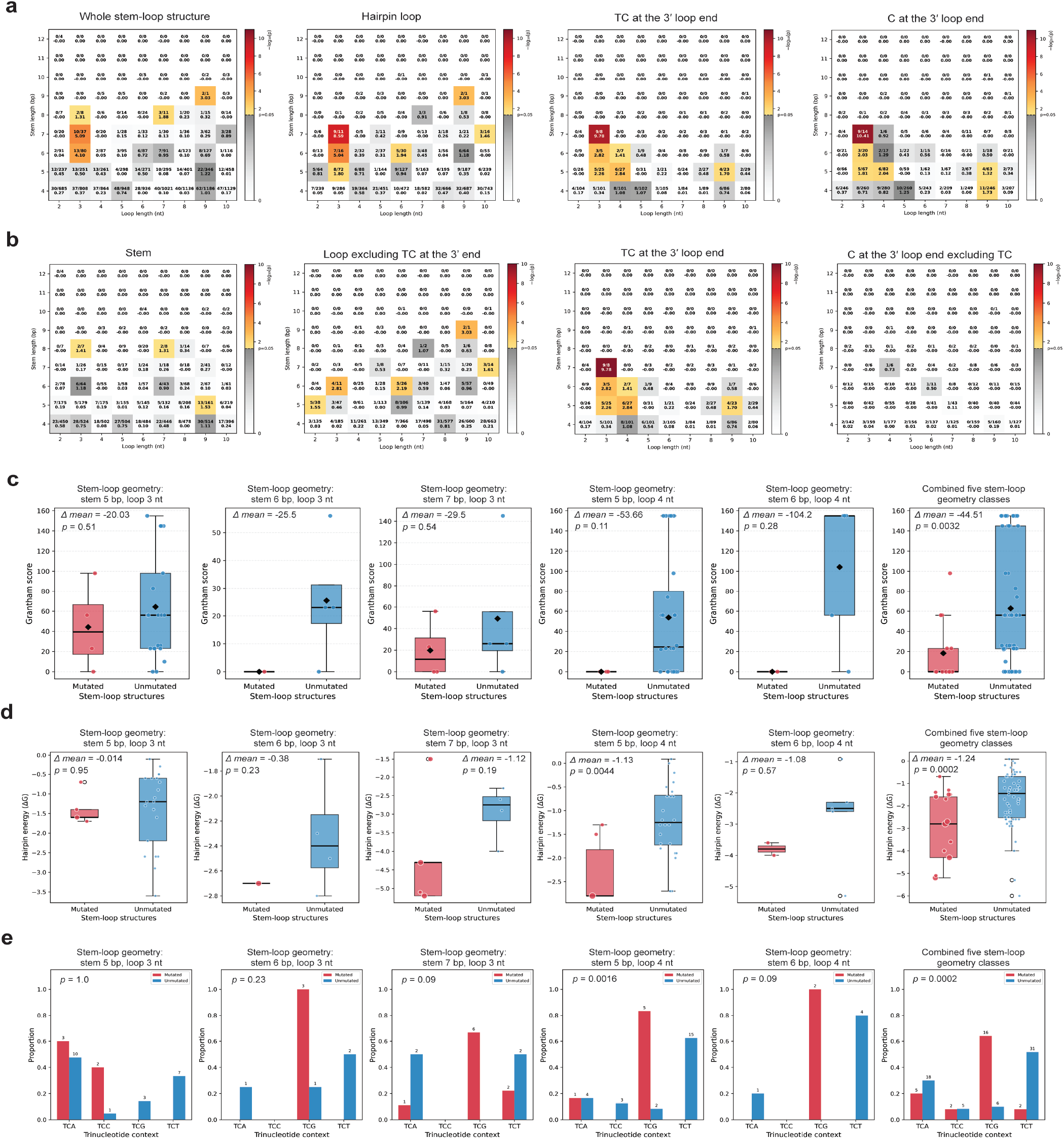
(a) APOBEC-signature mutation enrichment across geometries of stem-loop structures defined by stem and loop length. Stem-loop structures were predicted with no mismatches allowed in the stem, and all predicted structures were included without resolving overlaps. Heatmaps show the statistical significance of enrichment for different hairpin/ cruciform-associated structural contexts. In addition to the p-value, each cell displays the number of mutations and the number of potential target sites, separated by a slash. Gray shading indicates non-significant enrichment (*p* ≥ 0.05). (b) APOBEC-signature mutation enrichment across geometries of stem-loop structures, shown separately for distinct structural and motif-defined regions of predicted structures: stem positions, loop positions excluding TC motifs at the 3′ end of the loop, TC motifs at the 3′ end of the loop, and cytosines at the 3′ end of the loop excluding TC motifs. In addition to the p-value, each cell displays the number of mutations and the number of potential target sites, separated by a slash. (c) Comparison of Grantham scores, estimating the predicted physicochemical impact of amino acid substitutions, between observed APOBEC-signature C→T substitutions and hypothetical C→T substitutions at non-mutated TC motifs located at the 3′ end of the hairpin loop within the five significantly enriched classes of stem-loop structures defined by specific stem and loop lengths. (d) Comparison of hairpin folding free energy (Δ*G*) in single-stranded DNA between structures carrying APOBEC-signature substitutions and non-mutated structures within the five significantly enriched classes of stem-loop structures defined by specific stem and loop lengths. (e) Comparison of the frequencies of the nucleotide immediately downstream of the TC motif between structures carrying APOBEC-signature substitutions and non-mutated structures within the five significantly enriched classes of stem-loop structures defined by specific stem and loop lengths.

To identify which structural elements contribute to the observed enrichment, we analyzed the components of stem-loop structures separately, including stem regions, loop positions excluding 3’-terminal C and TC motifs, TC motifs at the 3’ ends of hairpin loops, and cytosines at the 3’ ends of hairpin loops whose preceding nucleotide was not T (Fig. 4b, Fig. S8). Enrichment was detected predominantly for TC motifs located at the 3’ ends of hairpin loops, whereas some cases of significant enrichment for particular geometries, primarily with long stems, was observed in the stem or in other loop positions.

To account for multiple testing across stem length (4-12 bp), loop length (2-10 nt), structural contexts, and structure-selection strategies (1,620 tests in total), we performed an empirical global test using randomized genomic positions (see Methods). Although individual tests may produce occasional significant results by chance, the observed number of significant geometries greatly exceeded the null expectation. Thus, in randomized controls only two significant geometries were observed on average (Fig. S9b), compared with 45 in the real genome, suggesting that the expected false-positive fraction (FDR) among nominally significant geometries is below 5%.

### Hairpin Stability, Nucleotide Context at the +1 Positions Relative to TC, and Substitution Deleteriousness Influence APOBEC-Induced Mutation Frequency in IRs

For further analysis, we selected five groups of structures defined by their stem-loop geometry (i.e., specific stem and loop lengths) that exhibited the most significant enrichment of APOBEC-signature substitutions: three groups with the loop length of 3 nt and the stem lengths of 7, 6, and 5 bp, and two groups with the loop length of 4 nt and the stem lengths of 6 and 5 bp, respectively. Although these groups of structures were strongly enriched for APOBEC-signature substitutions, many structures within them remained un-mutated, prompting us to investigate factors that may influence the structure mutability.

We first hypothesized that unmutated structures correspond to genomic positions where APOBEC-signature substitutions would result in deleterious amino acid changes. To test this hypothesis, we calculated the Grantham score [13], which quantifies the physicochemical differences between amino acid pairs, for observed APOBEC-signature substitutions and for hypothetical C→T substitutions at non-mutated TC motifs located at the 3’ ends of the hairpin loop (Fig. 4c). Across all five structure groups, hypothetical substitutions in non-mutated structures exhibited higher Grantham scores than the observed substitutions in mutated structures, and the difference was statistically significant when all five groups of structures were analyzed together (permutation test, *p* = 3.2 × 10-3). These results suggest that the absence of mutations in these structures may reflect stronger negative selection, whereby substitutions with greater functional impact are less likely to become fixed.

We next examined whether secondary-structure stability was associated with mutation occurrence by evaluating both the free energy of cruciform formation and the folding free energy of hairpins formed in single-stranded DNA. Both measures differed significantly between mutated and non-mutated structures, but the associations were of opposite direction. The difference in single-stranded DNA hairpin folding energy had stronger statistical support, with mutated hairpins exhibiting lower Δ*G* values and thus greater stability (permutation test, *p* = 2 × 10^-4^, Fig. 4d). In contrast, cruciform formation energies were significantly higher for mutated than for non-mutated structures (permutation test, *p* = 4.8 × 10^-3^, Fig. S9a). However, under the simplified energy model, the stem regions have equivalent energies in the hairpin and linear duplex conformations, so the calculated free energy change is largely determined by the energetic difference between the loop regions and the corresponding segments of the linear duplex. For 3-4 nt loops containing the TC motif, sequence variation is restricted to the remaining one or two nucleotides immediately upstream of TC. Consequently, this association may reflect APOBEC3 sequence specificity rather than an independent contribution of the cruciform stability.

Finally, we analyzed nucleotide frequencies at the +1 position immediately downstream of the target TC motif. Previous studies have shown that TCA is the predominant APOBEC mutation context in cancer genomes [9], whereas APOBEC-mediated editing of yeast transcripts and viral RNA preferentially occurs in the TCG context [14]. Consistent with the latter observations, TCG was the most frequent triplet context among APOBEC-signature substitutions in stem-loop structures and was significantly more frequent than the other TCN triplet contexts (permutation test, *p* = 2 × 10^-4^; Fig. 4e). We also examined the –1 position immediately upstream of the TC motif but detected no nucleotide preference in these groups of structures.

### Long-Stem, Short-Loop Stem-Loop Structures Carry Local Predictive Information for APOBEC-Signature Mutations, Whereas Guanine at the +2 and +3 Downstream of TC Contributes to Genome-Wide Prediction

To assess the extent to which the observed preference for stem-loop structures determines the localization of APOBEC-signature substitutions in the MPXV genome, we developed a set of predictive models incorporating hairpin/cruciform-forming propensity together with other genomic features, including local sequence context and the estimated deleteriousness of substitutions. We first assembled an additional dataset of MPXV genomes obtained from the GISAID database, comprising 2,301 sequences. Whole-genome sequence alignment (see Data Availability) identified 2,307 APOBEC-signature mutated sites corresponding to 3,127 independent mutational events. Together with the 2022 MPXV outbreak dataset (56 APOBEC-signature mutated sites) and the West African MPXV dataset (838 mutated sites corresponding to 1,089 independent events), this yielded a total of 4,272 APOBEC-signature mutational events occurring at 2,918 distinct genomic positions. Overall, these positions accounted for 11.8% of all TC/GA motifs in the MPXV genome. Based on the IRs predicted in the preceding analyses, we constructed a feature set describing TC motifs located at the 3′ ends of hairpin loops. Hairpin-related features included the presence and position of the loop-associated TC motif, stem and loop lengths, and the predicted folding free energy as a measure of structural stability. We additionally included the Grantham score as an estimate of the deleteriousness of the corresponding C→T substitution, as well as the nucleotides immediately flanking the TC motif (positions −1 and +1 relative to TC). These features were calculated for all TC/GA motifs in the MPXV genome and used as input variables for model training. The combined mutation data from the three datasets were used to define the response variable: TC/GA positions affected by an APOBEC-signature mutation were assigned to the positive class, whereas all remaining TC/ GA positions were assigned to the negative class.

We first evaluated whether the stem-loop structures contributes to the genome-wide prediction of APOBEC-signature mutations. For this analysis, we used all stem-loop structures predicted under the following parameters: minimum stem length of 4 bp, maximum stem length of 30 bp, maximum loop length of 20 nt, and no mismatches allowed within the stem. We trained eight classical machine-learning classifiers using two feature sets, either including or excluding the hairpin-related information (Fig. 5a). We used an equivalence margin of ±0.01 AUC, considering differences within this range practically negligible. Adding the stem-loop features did not significantly improve the performance of any classifier (paired bootstrap, 2000 resamples; see Methods). For four classifiers, the change in the AUC was within the equivalence margin. For three classifiers – random forest, XGBoost and naive Bayes – the performance was significantly worse (Holm-adjusted p = 0.004), potentially due to an increased number of features.

**Figure 5.**
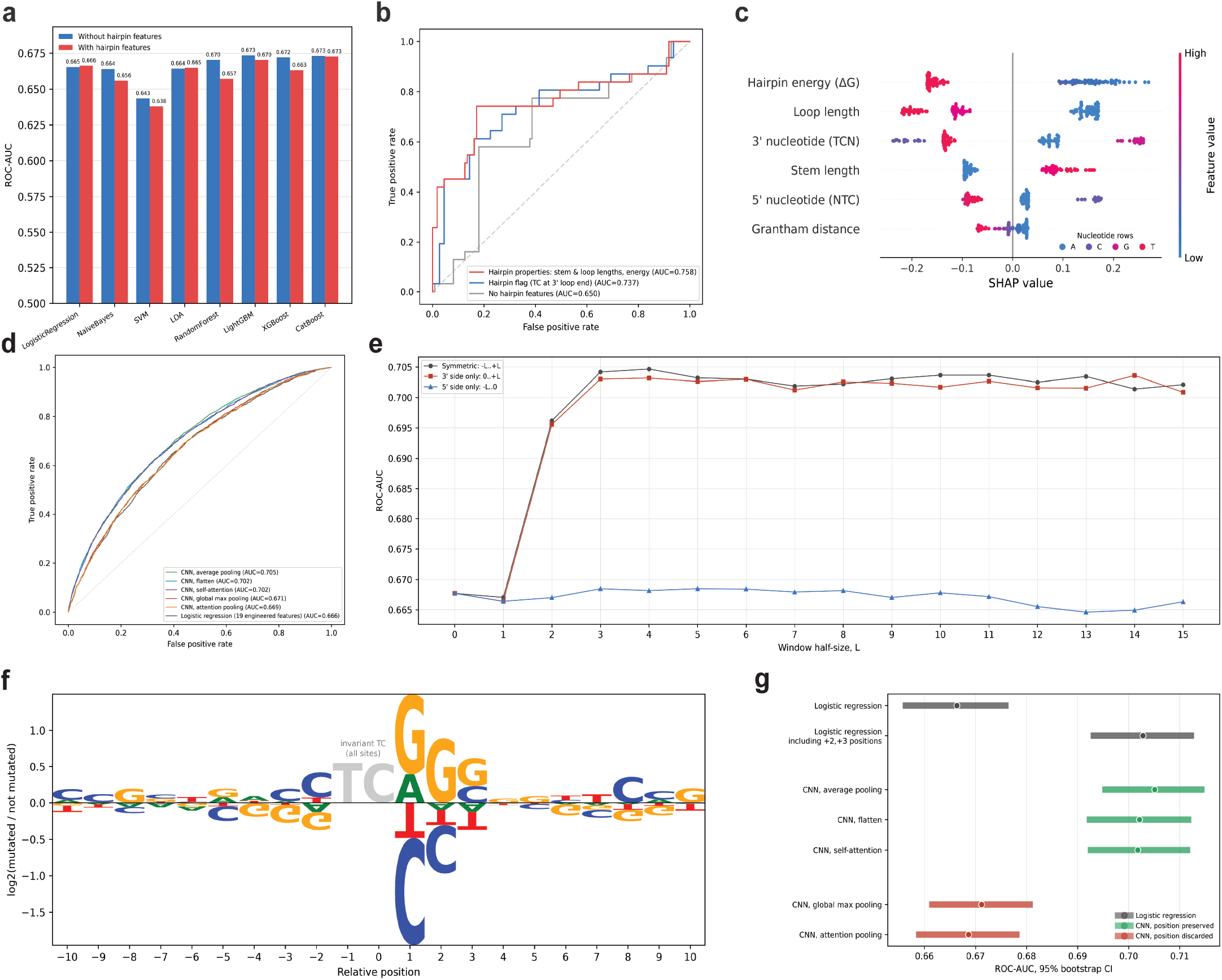
(a) Genome-wide ROC-AUC values for eight classical machine-learning models trained with or without hairpin/ cruciform-related features. (b) ROC curves for logistic regression models trained on a pair-matched dataset comprising TC/ GA sites at the 3′ ends of loops in stem-loop structures with long stems (5-7 bp) and short loops (3-4 nt), paired with matched sites outside structures. Models used three alternative feature sets: non-hairpin features alone, non-hairpin features combined with an indicator of hairpin presence, or non-hairpin features combined with detailed hairpin descriptors (stem length, loop length and folding free energy). (c) SHAP summary plot showing the contribution of individual features to logistic regression predictions. Features are ordered along the y-axis by decreasing mean absolute SHAP value, with the most influential features at the top. For numerical features, colors indicate feature values, ranging from low (blue) to high (red). For nucleotide features, colors indicate nucleotide identity: A (blue), C (purple), G (magenta), and T (red). (d) Genome-wide ROC curves for CNN models using different feature-aggregation strategies, including average pooling, global max pooling, flattening, attention pooling, and self-attention. Logistic regression is shown for comparison. (e) ROC-AUC of position-preserving CNN models as a function of sequence-window size. Windows were extended symmetrically around the TC motif or independently in the 3′ or 5′ direction. (f) Sequence-context enrichment around TC motifs, shown as the log2 ratio of nucleotide frequencies at mutated versus non-mutated sites. (g) Comparison of genome-wide ROC-AUC values and 95% bootstrap confidence intervals for logistic regression, logistic regression additionally including nucleotides at positions +2 and +3, and CNN architectures with position-preserving or position-discarding aggregation.

We also tested whether protein-level estimates of the substitution deleteriousness could improve prediction relative to the sequence-based Grantham score. To this end, we predicted three-dimensional structures of all MPXV proteins using AlphaFold [15] and applied two independent approaches, FoldX [16] and DDGun [17], to estimate mutation-induced changes in the protein stability. For seven of eight classifiers, replacing the Grantham score with any of three stability estimates yielded statistically equivalent predictive performance within a margin of ±0.01 ROC-AUC (paired bootstrap, 2,000 resamples; Figs. S9c, d). Random forest was the sole exception, with all three stability estimates yielding lower predictive performance than the Grantham score.

We next assessed whether hairpin/cruciform-related features contribute to local prediction of APOBEC-signature mutations, focusing on the preferentially targeted stem-loop structure classes identified above, namely those with stem lengths of 5-7 bp and loop lengths of 3-4 nt. Specifically, we examined TC motifs located at the 3′ ends of loops in these hairpins, identifying 71 such positions in the MPXV genome. For comparison, each of these positions was pair-matched to a TC motif located outside a hairpin with the same values of the non-structural features, including the −1 and +1 nucleotide context and the Grantham score, resulting in a balanced dataset of 142 positions. On this dataset, logistic regression achieved a ROC-AUC of 0.76 (Fig. 5b), which was 0.09 higher than 0.67 achieved by the best-performing model on the full dataset (Fig. 5a). To determine how different levels of structural information contributed to local prediction, we compared three progressively expanded feature sets: (i) non-hairpin features only, (ii) non-hairpin features plus an indicator of the hairpin presence, and (iii) non-hairpin features plus detailed descriptors of the hairpin geometry and stability, including folding free energy, stem length, and loop length. The performance improved progressively as more detailed structural information was incorporated (Fig. 5b), supporting a moderate local predictive signal associated with the presence and properties of these long-stem, short-loop hairpins.

We performed SHAP (SHapley Additive exPlanations) feature-importance analysis [18] over 100 iterations, each using the same 71 hairpin-associated positions and a random sample of 71 TC/GA positions (see Methods). The hairpin folding free energy made the largest contribution to prediction (Fig. 5c). The loop length and the 3′ nucleotide ranked next, followed by the stem length and the 5′ nucleotide, whereas the Grantham score had the smallest contribution.

We further asked whether additional information contained in the nucleotide context surrounding TC motifs could improve the genome-wide prediction. To capture sequence features beyond the immediately flanking nucleotides, we trained convolutional neural network (CNN) models using different feature-aggregation strategies, including pooling and attention-based mechanisms. CNN architectures that preserved positional information consistently outperformed models using position-invariant aggregation (Fig. 5d), suggesting that specific nucleotide positions relative to the TC motif contribute differently to the mutation probability. To identify positions responsible for this improvement, we trained position-preserving CNNs using progressively increasing sequence windows around the TC motif. We varied the size of the complete window as well as its upstream and down-stream portions independently. The prediction performance increased markedly as the downstream window was extended from +1 to +3 nt and then reached a plateau, whereas extension of the upstream window produced little additional improvement (Fig. 5e). These results indicated that nucleotides at positions +2 and +3 downstream of TC contain additional information associated with APOBEC-signature mutation frequency. Analysis of nucleotide composition at these positions revealed enrichment of guanine at both +2 and +3, extending the preference for guanine at the +1 position described in the preceding subsection (Fig. 5f). Consistent with this result, adding the identities of the +2 and +3 nucleotides as explicit features to the logistic regression model increased its predictive performance to a level comparable to that of the position-preserving CNNs (Fig. 5g). The same sequence logo (Fig. 5f) also showed a predominance of cytosine and thymine (Y) immediately upstream of the TC motif. This pyrimidine preference persisted when the analysis was restricted to TCA motifs (Fig. S9e), consistent with the YTCA sequence context characteristic of the APOBEC3A-mediated mutagenesis. We previously observed this preference in a smaller dataset [19], but it contrasts with the findings of an experimental study [20]. Together, these analyses indicate that the extended YTCGGG sequence context is preferentially associated with APOBEC-signature mutations in the MPXV genome.

## Discussion

This study began with a reevaluation of the report that APOBEC-induced mutations in the MPXV genome preferentially occur in stem-loop structures and their surrounding regions. Our reanalysis indicated that the original dataset, comprising more than 50 mutations from the 2022 outbreak, was insufficient for deriving statistically robust conclusions. To resolve this ambiguity, we reevaluated APOBEC preferences for stem-loop structures using a larger dataset of MPXV genomes from West Africa. Our analysis of this dataset shows that APOBEC-like mutations in the MPXV genome are enriched at TC motifs located at the 3′ ends of hairpin loops, with no enrichment at TC motifs in other regions of the loops, within stems, or near hairpin/cruciform structures. Similar preferences have been reported in genomes of cancer types with known APOBEC activity [5]. We also observed greater enrichment in stem-loop structures with longer stems and shorter loops, with the strongest enrichment in structures with three-nucleotide loops and the maximum stem length allowed by the genomic sequence under consideration. This tendency was previously described for APOBEC-induced mutations in cancer genomes by Buisson and colleagues [6], who showed that these mutations occur more frequently at TC motifs located at the 3′ ends of hairpin loops and that the mutation frequency increases with the stem length. They also found that hairpins with three-nucleotide loops exhibit the highest mutability, with mutation frequency gradually decreasing as the loop length increases. Hairpins with loops shorter than three nucleotides, however, did not appear to be preferred substrates in either our analysis or the previous study [5], possibly due to the instability of these structures.

Among groups of stem-loop structures defined by identical stem and loop lengths, the strongest preference for TC motifs at the 3′ ends of hairpin loops was observed in five structural classes, comprising three-nucleotide loops with stems of 5-7 bp and four-nucleotide loops with stems of 5-6 bp. The MPXV genome contains no longer inverted repeats with three-nucleotide spacers and only one longer inverted repeat with a four-nucleotide spacer. Although these groups of stem-loop structures were enriched for mutations in the MPXV genome, not all individual structures within them contained mutations. Given that purifying selection is expected to eliminate deleterious substitutions, we assessed whether the absence of mutations in individual structures within these groups was associated with the predicted deleteriousness of potential TC→TT substitutions. The observed substitutions in mutated stem-loop structures had significantly lower predicted deleteriousness scores than hypothetical substitutions in non-mutated structures. To examine structural stability as a potential factor influencing mutability, we assessed both types of secondary structures that can form at inverted repeats: cruciforms in double-stranded DNA and hairpins in single-stranded DNA. Mutated and non-mutated structures differed significantly under both models, but the direction of the association was opposite: mutated stem-loop structures were predicted to be more stable as single-stranded hairpins and less thermodynamically favorable as cruciforms. The stronger association observed for hairpins formed in single-stranded DNA suggests that these structures in particular may contribute to the APOBEC-mediated mutagenesis. Analysis of nucleotide frequencies at the +1 position immediately downstream of TC further showed that mutated structures were enriched for the TCG motif, whereas the other TCN contexts showed no statistically significant differences between mutated and non-mutated groups. Recently, TCG has been reported as the predominant context for APOBEC-mediated mutations in viral RNA and yeast transcripts [14].

We also investigated the extent to which stem-loop structures and the other aforementioned factors determine the distribution of mutations across all TC/GA motifs in the MPXV genome. To this end, we developed and evaluated several machine learning models, including convolutional neural networks, for predicting the genomic locations of candidate APOBEC-induced mutations in the MPXV. Hairpin/cruciform-related features provided a moderate local predictive signal for TC motifs associated with long-stem, short-loop structures. However, incorporating these features did not improve the genome-wide prediction. At the genome-wide level, our analyses highlighted the importance of the nucleotide immediately upstream of TC and three nucleotides downstream. Pyrimidines predominated at the upstream position, while guanine was enriched at all three downstream positions, supporting a preference for the extended YTCGGG sequence context. Adding the downstream nucleotides at +2 and +3 to logistic regression brought its performance to a level comparable to that of position-preserving CNNs, suggesting that much of the additional predictive information was captured by this short, position-specific sequence context.

Overall, our study shows that candidate APO-BEC-induced mutations accumulating in the MPXV genome following zoonotic transmission are non-randomly distributed and are associated with the nucleotide context and predicted DNA secondary structures. Stem-loop structures can be predicted bioinformatically, while complementary approaches can estimate their stability and the deleteriousness of substitutions in protein-coding regions. Deleterious substitutions are less likely to be observed because they tend to be removed by purifying selection. Combined with the analysis of nucleotide context, these methods can help explain, and to a limited extent predict, where substitutions accumulate. Together, these analyses provide a framework for understanding how human APOBEC3-mediated antiviral activity may shape viral genome evolution, with potential implications for vaccine design.

## Methods

### MPXV genomes and sequence alignment

The MPXV reference genome (accession GCF_014621545) was downloaded from GenBank [21]. The first dataset, containing substitution positions identified during the 2022 MPXV outbreak, was obtained from [4]. Genomes for the second dataset were obtained from [8] and aligned using MAFFT [22]. For the third dataset, 2,301 MPXV genomes deposited between 01.01.2022 and 30.01.2026 were downloaded from GISAID [23]. To reduce the computational cost of multiple sequence alignment, the genomes were grouped by sequence similarity, aligned within groups and subsequently merged into a single alignment. Pairwise genomic similarities were calculated using fastANI [24] with default parameters, except that --fragLen was reduced from 3,000 to 1,000 bp. The resulting genome distance matrix was used for complete-linkage hierarchical clustering implemented in SciPy. The clustering threshold was adjusted to obtain approximately 200-300 genomes per cluster. Genomes within each cluster were aligned using MAFFT with the parameters --maxiterate 1000, --retree 5, --kimura 1, and --adjustdirection. The resulting cluster alignments were merged into a single alignment using the MAFFT –merge option.

### Prediction hairpin/cruciform structures

Inverted repeats (IRs) were detected with Palindrom Analyzer [12] and EMBOSS package [11]. Each IR was represented by its outer genomic span (from the earliest arm start to the latest arm end). Based on initial IR prediction the interval-overlap graph was constructed. In this graph nodes are IRs and an edge connects two IRs whose spans overlap by ≥1 bp. Overlap groups were defined as the connected components of this graph.

Within each overlap group, four strategies were applied: (1) select the single structure with the lowest predicted free energy (as reported by PalindromAnalyzer output) as the most-stable representative; (2) apply a greedy non-overlapping procedure that iteratively selects the most stable structure and removes all overlapping candidates until none remain; (3) use dynamic programming to choose structures (or hairpin loops, depending on the experiment) that maximize total genomic coverage within the group; and (4) select a single representative that minimizes genomic coverage by the structure itself (or by its loop, as appropriate). For binomial approximations, the probability of success parameter *0*– was defined as the fraction of the genome covered by the selected structures (or loops), i.e., total covered base pairs divided by the genome length.

### Hairpin and cruciform energy calculations

For hairpins inferred from EMBOSS, free energies were calculated using a nearest-neighbor (NN) model [25], following the approach used in Palindrome Analyzer. In this model, helical stability depends on interactions between adjacent base pairs, with the total free energy calculated by summing the corresponding contributions. The free energies of the linear duplex and hairpin conformations were calculated separately, with the hairpin calculation incorporating terms accounting for mismatches and loop length. The free energy change associated with cruciform formation was calculated as Δ*G* = 2*G*_*hairpin*_*-G*_*linear*_, reflecting the formation of two opposing hairpins, one on each strand. This quantity was used as a proxy for cruciform stability relative to the linear duplex. Thermodynamic parameters and equations were obtained from the DNA section of the Nearest Neighbor Database (https://rna.urmc.rochester.edu/NNDB/).

### Independent mutational events calculation

To avoid counting the same mutation multiple times across downstream branches, mutations were mapped as independent events on the phylogenetic tree. A multiple-sequence alignment of MPXV genomes was used to reconstruct a maximum-likelihood phylogeny in IQTREE [26], with the substitution model selected automatically using the --substitution-model AUTO option. Then, the resulting tree was reconstructed using Augur refine (timetree) into a time-resolved phylogeny [27]. The NC_063383.1 reference genome was kept as the root and date inference was performed using marginal maximum likelihood estimation under a constant coalescent model. Augur’s covariance-aware mode was disabled using the --no-covariance flag. Finally, ancestral nucleotide sequences for each internal node and branch-specific substitutions were jointly inferred by maximum likelihood with Augur ancestral.

### Permutation-based estimation of the false discovery rate

The false discovery rate (FDR) of tests for APOBEC3 mutation enrichment in groups of stem-loop structures defined by stem and loop lengths was estimated using genome sequence permutations. In each of 1,000 iterations, the MPXV reference sequence was randomly permuted to preserve its mononucleotide composition while disrupting its native sequence organization. The observed mutation positions were held fixed. Each permuted genome was analyzed using the same pipeline as the original genome. Each iteration comprised 1,620 tests (5 selection strategies × 4 structural categories × 9 stem lengths × 9 loop lengths), and the number yielding *p* < 0.05 was recorded. The empirical FDR was estimated as the mean number of significant tests across the 1,000 permuted genomes divided by the number of significant tests for the original genome.

### Classic machine learning models

Each of the 24,695 candidate APOBEC3 target sites (TC/GA) of the MPXV reference genome (NC_063383.1) was represented as one observation. The binary outcome indicated whether the site was mutated in at least one of the three mutation datasets (2,918 positive sites, 11.8%). Sequence-based predictors included the Grantham score for the corresponding C→T substitution and the two bases immediately 5′ and 3′ of the TC dinucleotide. Grantham scores were set to 0 for synonymous substitutions and intergenic sites, and to 215 for nonsense substitutions. Both flanking bases were one-hot encoded over A, C, G and T. Stem-loop structure predictors included an indicator of whether the site was located at the 3′ end of a hairpin loop, the corresponding loop and stem lengths, and three thermodynamic stability metrics. These metrics were the folding energy of the single-stranded DNA hairpin, the energy of the same region as a linear double-stranded duplex, and the free energy of cruciform extrusion. All three metrics were computed using a nearest-neighbor model and, as a consistency check, ViennaRNA [28]. For sites not located at the 3′ end of a hairpin loop, undefined hairpin metrics were set to 0. All features were standardized to zero mean and unit variance. To prevent information leakage, the mean and standard deviation of each feature were calculated using only the training data in each cross-validation fold and then used to standardize both the training and held-out data.

Classification was done using eight classical algorithms: logistic regression, Gaussian naive Bayes, a support vector machine with an RBF kernel, linear discriminant analysis, random forest, and the gradient-boosting implementations LightGBM, XGBoost and CatBoost. The gradient-boosting models were restricted to 100 trees of depth 3. All remaining hyperparameters were set to their library defaults, with a fixed random seed used throughout. Class imbalance was compensated by balanced class weighting: class_ weight=’balanced’ for the scikit-learn algorithms, scale_pos_weight for XGBoost, and balanced automatic class weights for CatBoost. Model training and evaluation used 10-fold stratified cross-validation, with feature scaling parameters and model parameters estimated exclusively from the training data within each fold. Out-of-fold predicted probabilities were pooled across folds to calculate the area under the receiver operating characteristic curve (ROC-AUC). Differences in ROC-AUC between models were assessed using a paired bootstrap with 2,000 resamples of sites drawn with replacement. Both models were evaluated on the same sites within each resample, and the difference in ROC-AUC was calculated, preserving the dependence between their performance estimates. The resulting bootstrap distribution was used to derive a 95% percentile confidence interval. Two-sided bootstrap p-values were calculated as twice the smaller of the numbers of resampled differences ≤0 and ≥0, divided by the total number of resamples, with a lower bound of 1/2,000. P-values were adjusted for multiple comparisons across the eight algorithms using the Holm procedure. Statistical equivalence was assessed separately using a margin of ±0.01 ROC-AUC, representing the largest difference considered practically negligible. Equivalence was established only when the entire 95% confidence interval lay within this margin. Analyses were performed in Python with scikit-learn 1.9.0 [29], LightGBM 4.6.0 [30], XGBoost 3.1.3 [31] and Cat-Boost 1.2.10 [32].

### Deep learning models

The convolutional models combined two inputs: a nucleotide sequence window extending ±15 nt around the edited cytosine (31 positions) and the engineered feature vector used by the classical models. Sequences were one-hot encoded over A, C, G and T, with undetermined bases represented by all-zero columns. Engineered features were standardized using parameters estimated exclusively from the training fold. The sequence branch comprised a single one-dimensional convolutional layer with eight filters of width 5, zero padding and a ReLU activation, producing eight filter responses at each of the 31 positions. Five approaches were used to convert the feature map into a vector: (i) flattening, which preserved filter responses at every position (“CNN, flatten”); (ii) average pooling over adjacent pairs of positions followed by flattening, which retained positional information at half the original resolution (“CNN, average pooling”); (iii) global maximum pooling, which retained the maximum response of each filter across positions (“CNN, global max pooling”); (iv) attention pooling, which scored each position using a learned weighted sum of its filter responses, converted these scores into weights summing to one using softmax, and used these weights to average the filter responses across positions (“CNN, attention pooling”); and (v) self-attention followed by average pooling and flattening, which integrated information across positions using a single multi-head scaled dot-product self-attention block with two attention heads, a residual connection and layer normalization (“CNN, self-attention”). For all five approaches, the resulting sequence vector was concatenated with the engineered features and passed to a fully connected prediction head comprising one hidden layer of 32 units with a ReLU activation and a single-logit output. Dropout at a rate of 0.3 was applied before and within the prediction head. For the architectural comparison, the flattening, average-pooling and self-attention models were grouped as “CNN, position preserved”, whereas the global maximum-pooling and attention-pooling models were grouped as “CNN, position discarded”.

Models were trained using the Adam optimizer (learning rate 10^-3^, weight decay 10^-4^) for 20 epochs with a mini-batch size of 256 by minimizing binary cross-entropy loss. To account for class imbalance, positive-site contributions to the loss were weighted by the ratio of negative to positive sites in the training fold, consistent with the weighting strategy used for the classical models. Performance was evaluated using the same 10-fold stratified cross-validation scheme as for the classical models. Feature standardization parameters and network parameters were estimated exclusively from the training data within each fold. Out-of-fold predicted probabilities were pooled across folds to calculate ROC-AUC. Random seeds for weight initialization and batch shuffling were fixed for each fold. To localise the sequence signal, the convolutional model with feature-map flattening (“CNN, flatten”) was re-trained with progressively larger sequence windows extending up to 15 nt from the target cytosine. Models were implemented in PyTorch 2.9.1 [33].

### SHAP-based analysis of local feature contributions

To quantify feature contributions locally, at the scale of individual stem-loop structures, SHAP (SHapley Additive exPlanations) analysis [18] was performed on a set combining TC sites at the 3′ ends of hairpin loops whose stem-loop geometries are associated with the elevated mutation frequency reported above (loop length 3 with stem lengths 5-7, and loop length 4 with stem lengths 5-6; 71 sites in total) with an equal number of sites drawn at random and without replacement from the 22,424 TC/GA sites that lie outside stemloop structures. SHAP values for the fitted model’s predicted probabilities were computed using the exact explainer in the shap package (version 0.52.0), which enumerates all feature coalitions, with an interventional (independent) masker and the 142 analysed sites as the background dataset. The one-hot encoded 5′ and 3′ bases were represented as two categorical predictors during explanation and converted back to their original one-hot encoding within the prediction function, yielding a single contribution for each predictor at each site. Random sampling was repeated 100 times using independent random seeds, with the model refitted and SHAP values recomputed at each iteration. Features were ranked within each iteration by their mean absolute SHAP values, and the final ordering was determined by their mean ranks across iterations. The SHAP summary plot displays the iteration whose feature ranking was closest to the final ordering, as measured by Spearman’s footrule distance.

### Protein structure models and stability predictions

Structural models of all 175 MPXV proteins were generated using the AlphaFold web server [15] with default parameters. For each protein, only the first returned model (suffix “_0”) was retained. Protein stability changes were predicted for all APOBEC3 target sites at which a C→T substitution results in an amino acid substitution (15,187 of 24,695 sites). Synonymous, intergenic, nonsense and within-stop-codon sites were left unscored, as protein stability changes were not defined for these sites. Structural models were first processed using the FoldX [16] RepairPDB command to optimise side-chain rotamers and relieve residual steric strain. Each substitution was then introduced using BuildModel, which rebuilds the mutated and neighbouring side chains and calculates the change in folding free energy (ΔΔ*G*) relative to the wild-type model. Each substitution was modelled in five independent runs, with the mean ΔΔ*G* used as the stability estimate and the standard deviation across runs retained as a measure of numerical reproducibility. Independently, ΔΔ*G* was predicted with DDGun [17] in both its sequence-only and structure-based modes, the latter using the same AlphaFold models.

## Data availability

All data and scripts supporting this study, including sequence alignments, mutation datasets, protein models, and scripts for performing the analyses and generating the figures, are available in the GitHub repository at https://github.com/KazanovLab/APOBEC3-Mpox-Hairpins.

## Acknowledgements

This study was supported by Assignment FFRW-2024-0004 (to G.V.P., E.V.M., R.K.A., and M.D.K.) and, in part, by RSF grant #25-14-00491 (protein stability analysis, to D.N.I.). We acknowledge the School of Molecular and Theoretical Biology, where this study was initiated. We thank Irina Ponomareva for designing the figures and preparing the preprint layout.

