## Supplemental Figures for "Hairpin Architecture and Extended Sequence Context Shape APOBEC3 Mutagenesis in Monkeypox Virus at Different Genomic Scales"

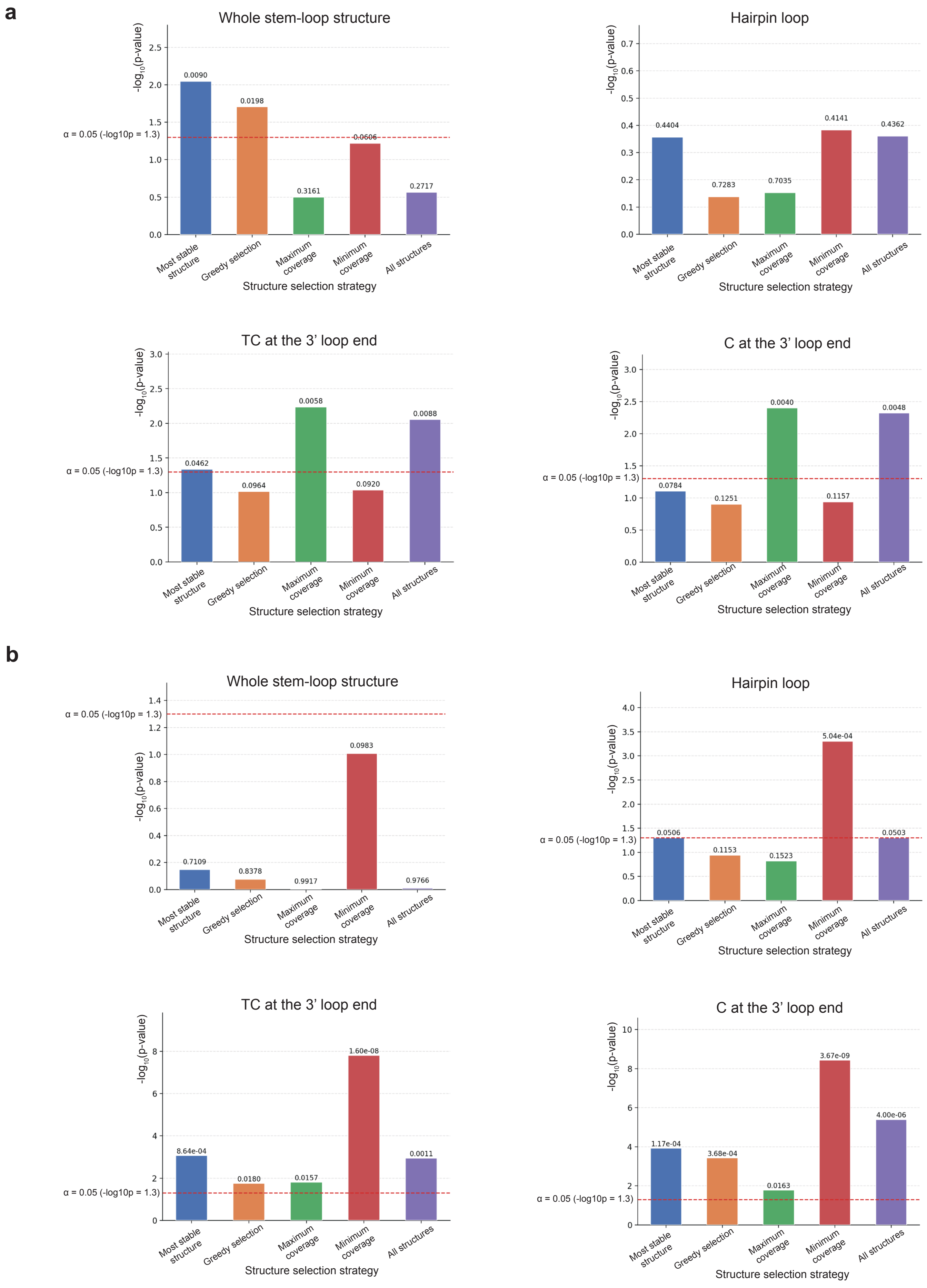


**Figure S1.** Statistical significance of enrichment analyses for APOBEC-signature mutational events in different hairpin/cruciform-associated structural contexts in West African and GISAID datasets. (a) Results for the West African dataset with inverted repeats predicted using PalindromeAnalyzer.(b) Results for the GISAID dataset with inverted repeats predicted using the EMBOSS palindrome program. For both panels, bars represent −log10(p) values and the dashed red line indicates the significance threshold (p = 0.05).

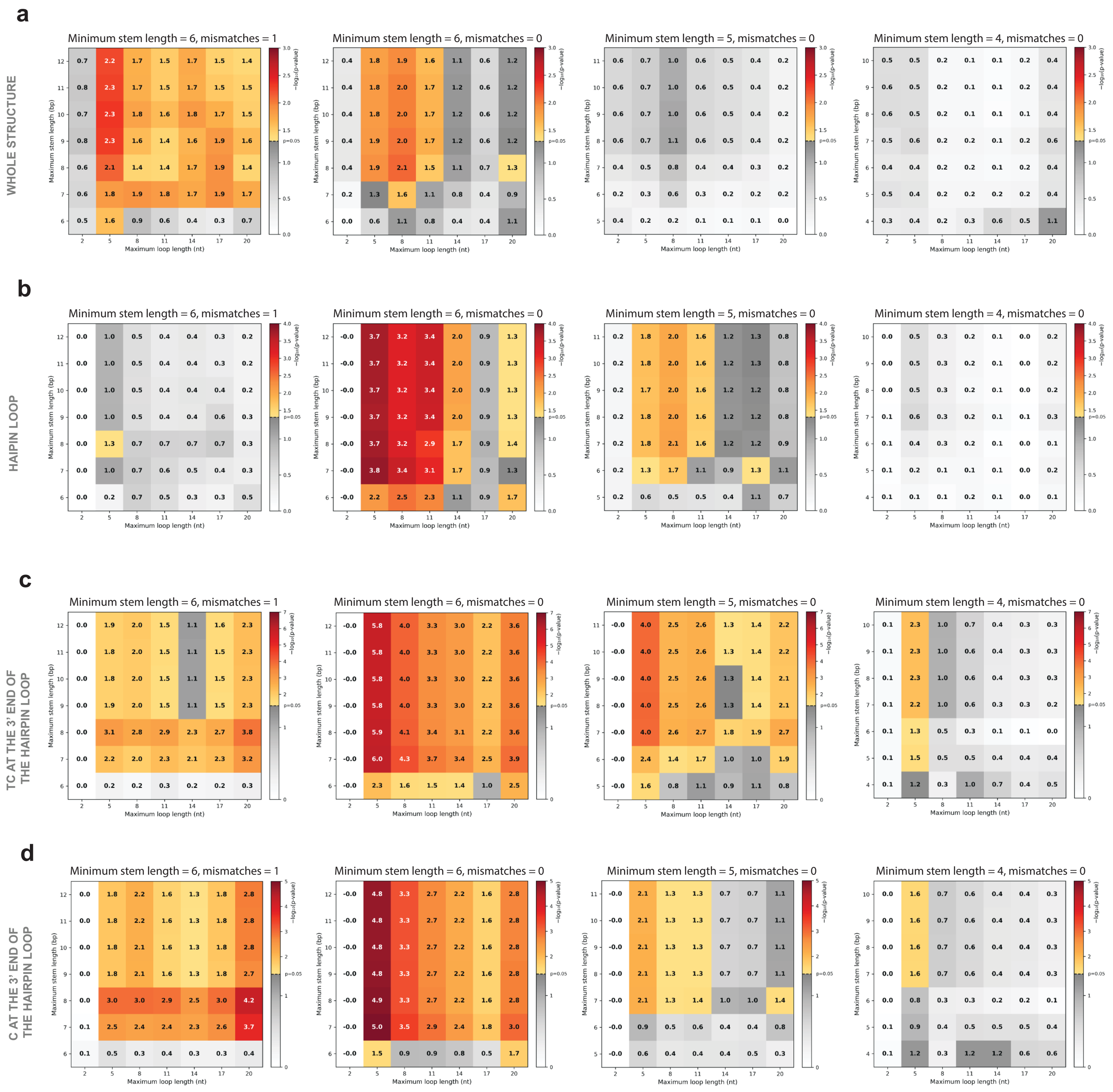


**Figure S2.** Statistical significance of enrichment analyses for APOBEC-signature mutational events in four hairpin/cruciform-associated structural contexts – (a) whole stem-loop structures, (b) hairpin loops, (c) TC motifs located at the 3′ ends of hairpin loops, and (d) cytosines located at the 3′ ends of hairpin loops – across different hairpin prediction parameters. Overlapping stem-loop structures were resolved by retaining the most stable structure within each overlapping group. Heatmaps show −log10(p) values for combinations of maximum stem length and maximum loop length. Analyses were performed with minimum stem lengths of 6 nt and 1 mismatch, 6 nt and 0 mismatches, 5 nt and 0 mismatches, and 4 nt and 0 mismatches. Gray shading indicates non-significant values ($p \geq 0.05$).

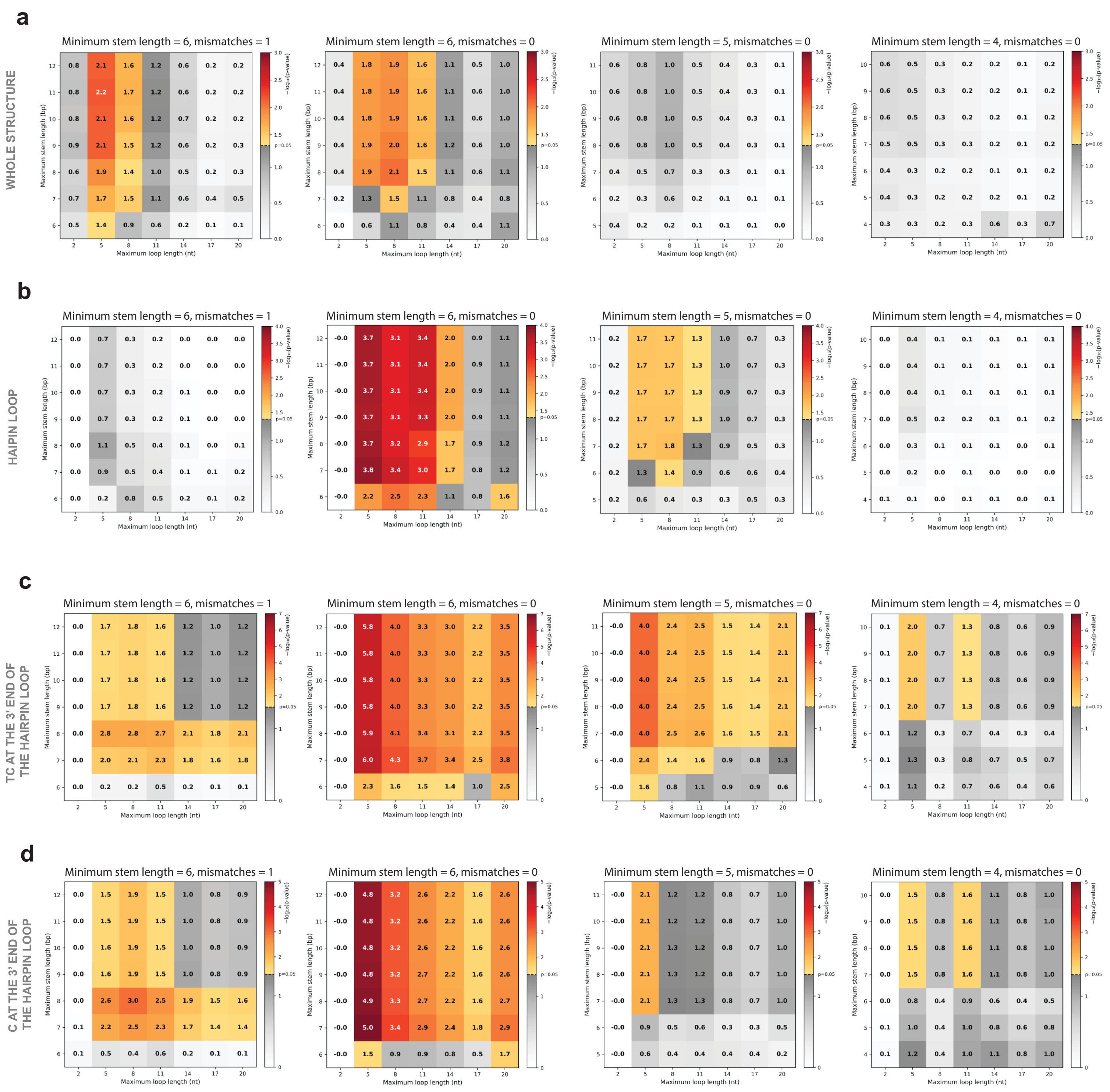


**Figure S3.** Statistical significance of enrichment analyses for APOBEC-signature mutational events in four hairpin/cruciform-associated structural contexts – (a) whole stem-loop structures, (b) hairpin loops, (c) TC motifs located at the 3′ ends of hairpin loops, and (d) cytosines located at the 3′ ends of hairpin loops – across different parameter settings for inverted repeat prediction. Overlapping stem-loop structures were resolved using a greedy selection strategy, iteratively selecting the most stable remaining structure and removing all overlapping structures until no overlaps remained. Heatmaps show −log10(p) values for combinations of maximum stem length and maximum loop length. Analyses were performed with minimum stem lengths of 6 nt and 1 mismatch, 6 nt and 0 mismatches, 5 nt and 0 mismatches, and 4 nt and 0 mismatches. Gray shading indicates non-significant values ($p \geq 0.05$).

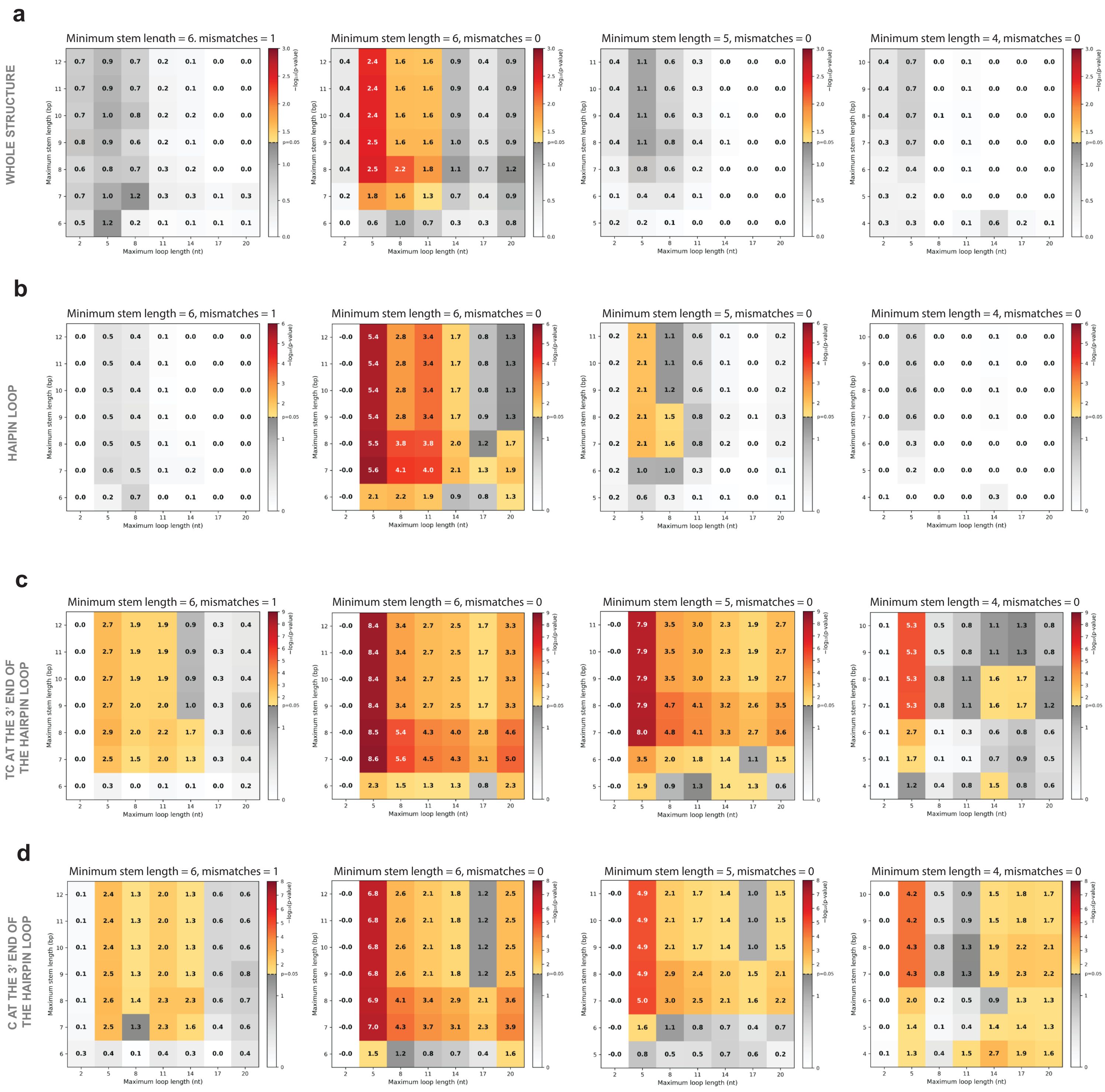


**Figure S4.** Statistical significance of enrichment analyses for APOBEC-signature mutational events in four hairpin/cruciform-associated structural contexts – (a) whole stem-loop structures, (b) hairpin loops, (c) TC motifs located at the 3′ ends of hairpin loops, and (d) cytosines located at the 3′ ends of hairpin loops – across different parameter settings for inverted repeat prediction. Overlapping stem-loop structures were resolved using a maximum coverage strategy, selecting a non-overlapping set of structures that maximized total genomic coverage. Heatmaps show −log10(p) values for combinations of maximum stem length and maximum loop length. Analyses were performed with minimum stem lengths of 6 nt and 1 mismatch, 6 nt and 0 mismatches, 5 nt and 0 mismatches, and 4 nt and 0 mismatches. Gray shading indicates non-significant values ($p \geq 0.05$).

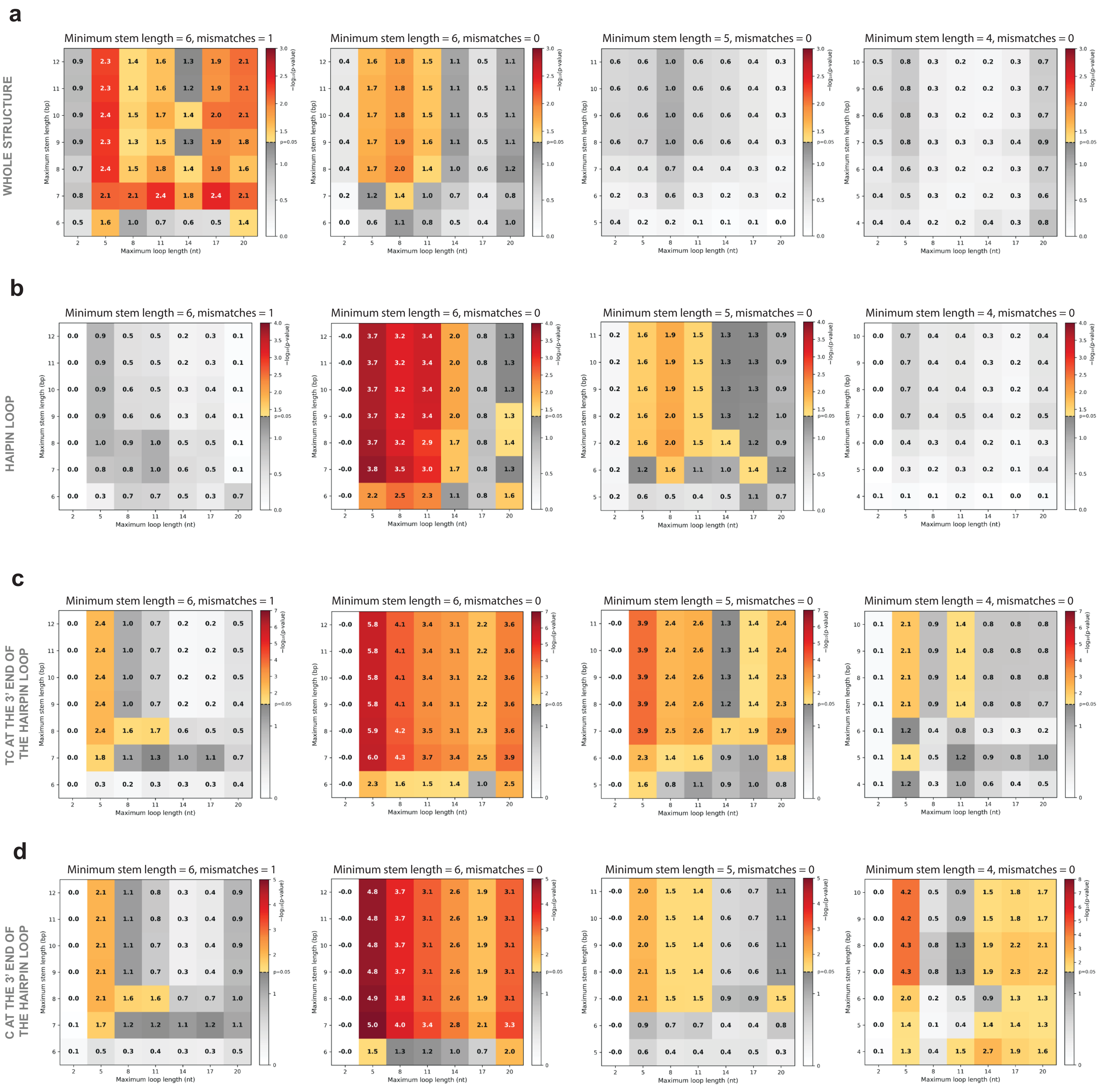


**Figure S5.** Statistical significance of enrichment analyses for APOBEC-signature mutational events in four hairpin/cruciform-associated structural contexts – (a) whole stem-loop structures, (b) hairpin loops, (c) TC motifs located at the 3′ ends of hairpin loops, and (d) cytosines located at the 3′ ends of hairpin loops – across different parameter settings for inverted repeat prediction. Overlapping stem-loop structures were resolved using a minimum coverage strategy, retaining the shortest structure within each overlapping group to minimize total genomic coverage. Heatmaps show −log10(p) values for combinations of maximum stem length and maximum loop length. Analyses were performed with minimum stem lengths of 6 nt and 1 mismatch, 6 nt and 0 mismatches, 5 nt and 0 mismatches, and 4 nt and 0 mismatches. Gray shading indicates non-significant values ($p \geq 0.05$).

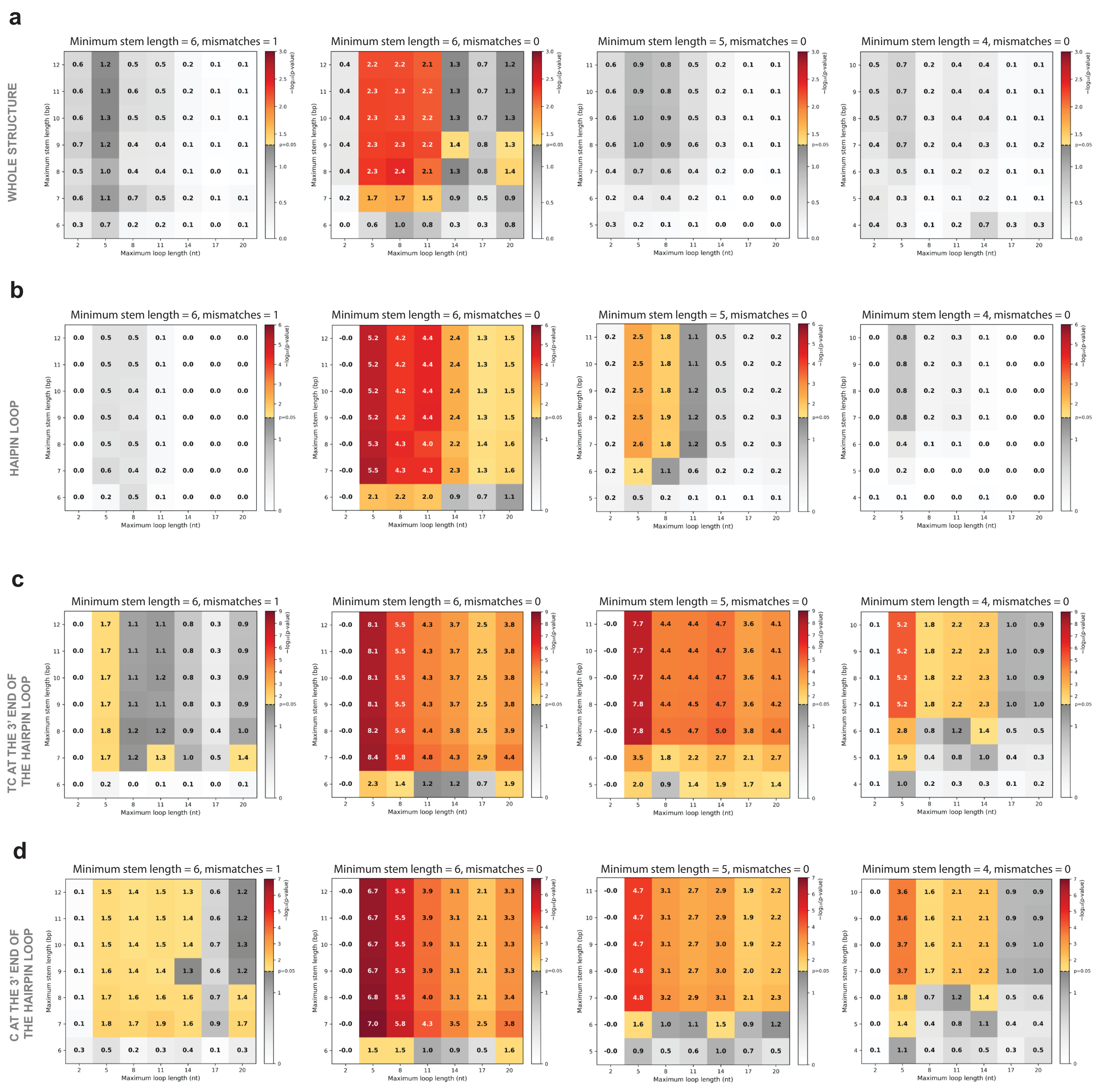


**Figure S6.** Statistical significance of enrichment analyses for APOBEC-signature mutational events in four hairpin/cruciform-associated structural contexts – (a) whole stem-loop structures, (b) hairpin loops, (c) TC motifs located at the 3′ ends of hairpin loops, and (d) cytosines located at the 3′ ends of hairpin loops – across different parameter settings for inverted repeat prediction. All predicted stem-loop structures were retained without applying any overlap resolution strategy. Heatmaps show −log10(p) values for combinations of maximum stem length and maximum loop length. Analyses were performed with minimum stem lengths of 6 nt and 1 mismatch, 6 nt and 0 mismatches, 5 nt and 0 mismatches, and 4 nt and 0 mismatches. Gray shading indicates non-significant values ($p \geq 0.05$).

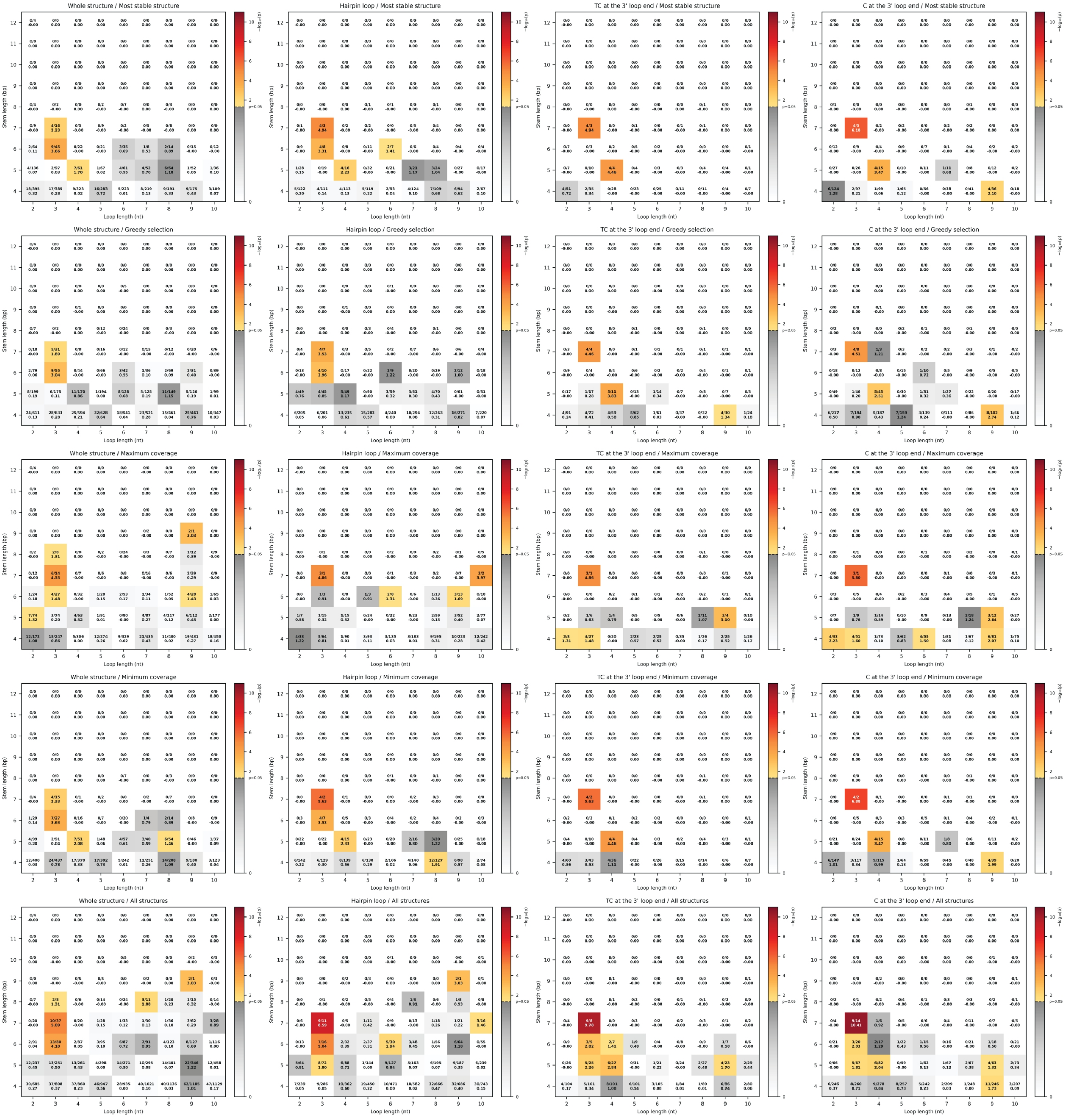

**Figure S7.** APOBEC-signature mutation enrichment across stem-loop structure geometries under different structural contexts and strategies for resolving overlapping structures. Inverted repeats were predicted with no mismatches allowed in the stem. Heatmaps show the statistical significance of APOBEC-signature mutation enrichment stratified by stem length and loop length. Gray shading indicates non-significant enrichment (p ≥ 0.05). In addition to the p-value, each cell displays the number of mutations and the number of potential target sites, separated by a slash. Columns correspond to four hairpin/cruciform-associated structural contexts, and rows correspond to five strategies for resolving overlapping structures.

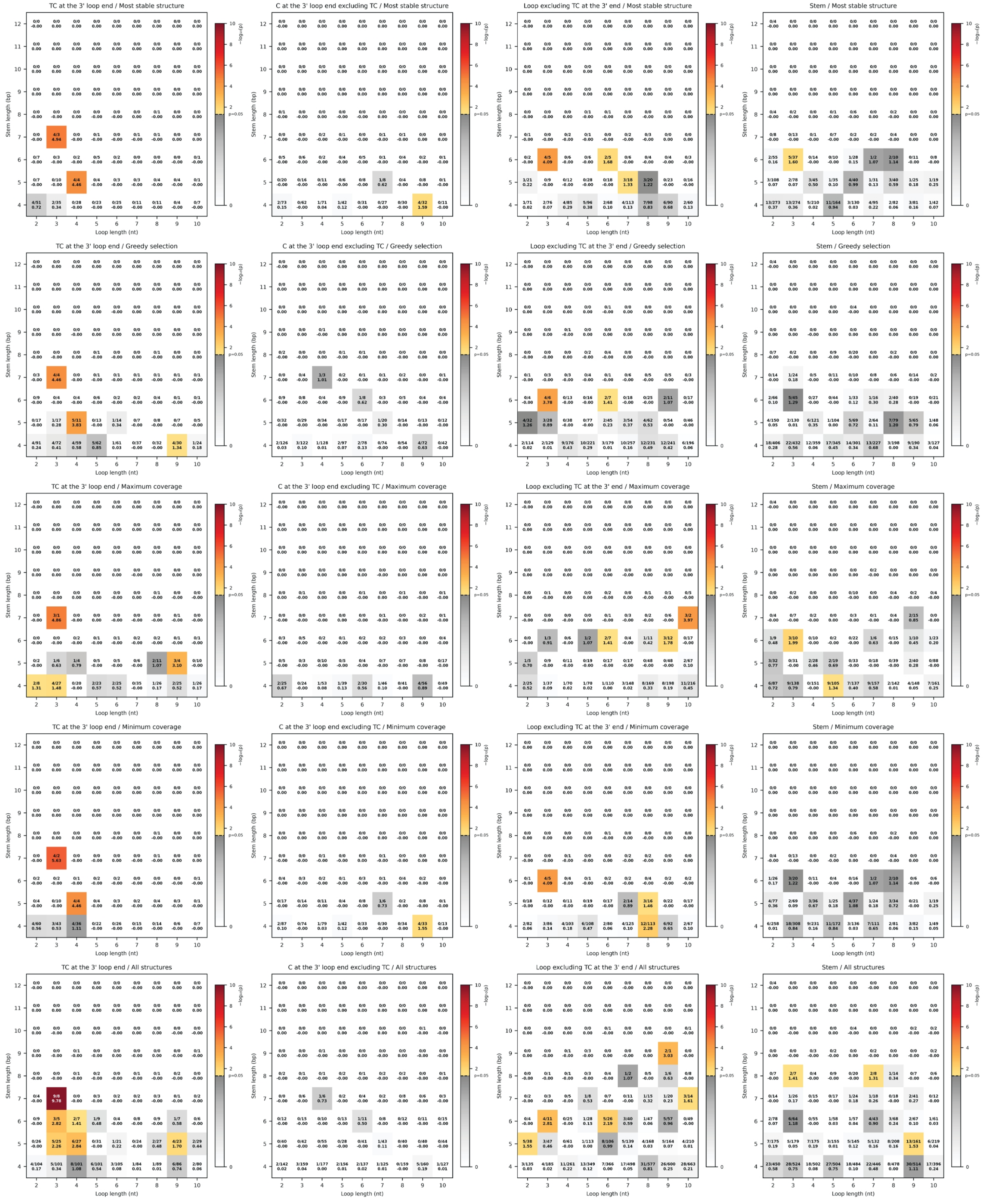


**Figure S8.** APOBEC-signature mutation enrichment across stem-loop structure geometries in distinct structural and motif-defined regions of predicted structures. Inverted repeats were predicted with no mismatches allowed in the stem. Heatmaps show the statistical significance of APOBEC-signature mutation enrichment stratified by stem length and loop length. Gray shading indicates non-significant enrichment (p ≥ 0.05). In addition to the p-value, each cell displays the number of mutations and the number of potential target sites, separated by a slash. Columns correspond to the analyzed regions: stem positions, loop positions excluding TC motifs at the 3′ end of the loop, TC motifs at the 3′ end of the loop, and cytosines at the 3′ end of the loop excluding TC motifs. Rows correspond to five strategies for resolving overlapping structures.

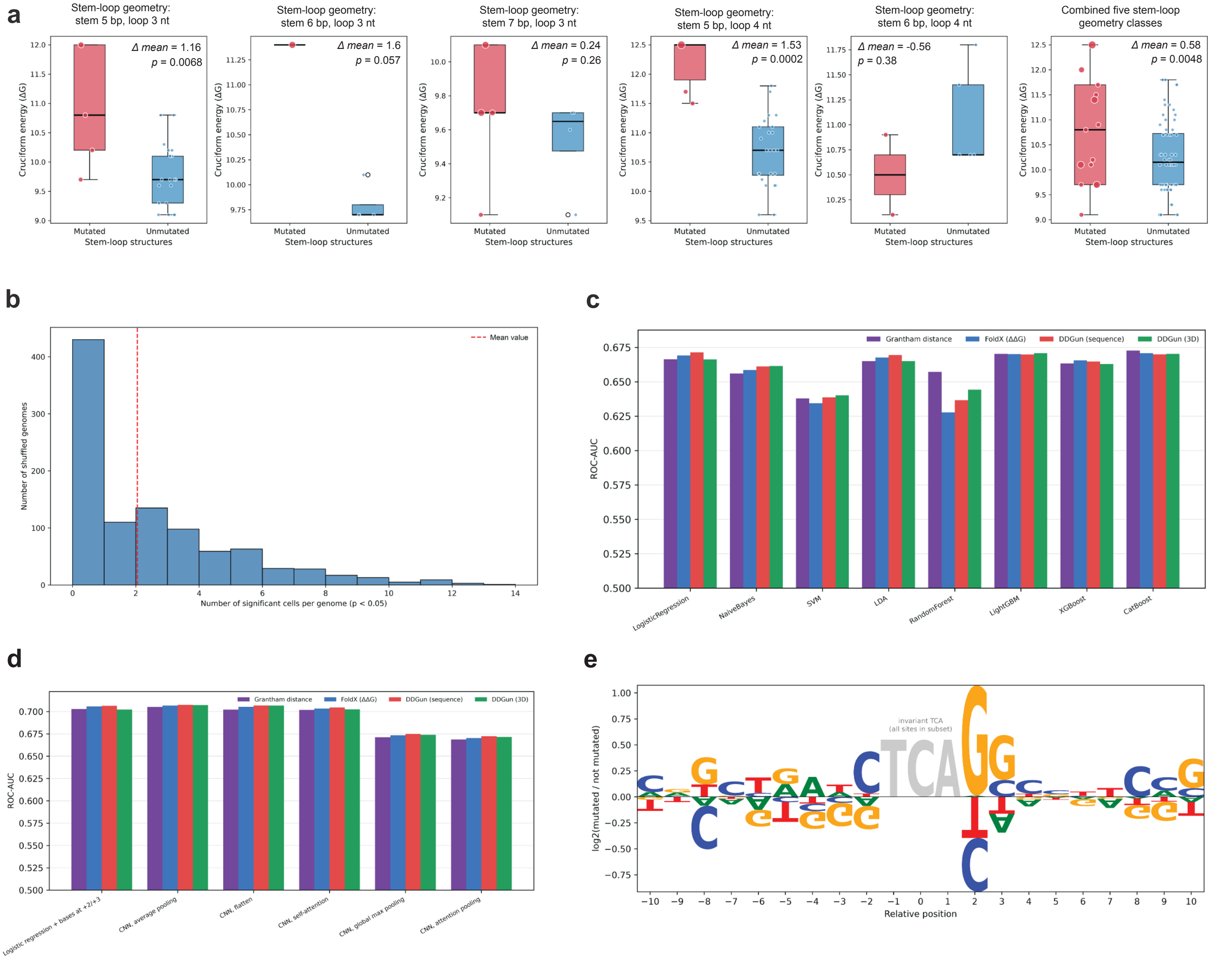


**Figure S9.** (a) Comparison of the free energy change associated with cruciform formation (ΔG) in double-stranded DNA between inverted repeat regions carrying APOBEC-signature substitutions and non-mutated regions within the five significantly enriched stem-loop structure classes defined by specific stem and loop lengths. (b) Empirical null distribution of the number of stem–loop structure geometry classes showing significant enrichment. APOBEC-signature mutation enrichment was assessed in randomized control genomes using the same analysis pipeline as for the real MPXV genome, including all tested stem lengths (4–12 bp), loop lengths (2–10 nt), structural contexts, and stem-loop structure selection strategies (1,620 tests in total). The histogram shows the number of geometry classes of stem-loop structures exhibiting nominally significant enrichment ($p < 0.05$) in each of 1,000 randomized genomes. The dashed red line indicates the mean number of significant geometry classes across randomized genomes. (c,d) Comparison of ROC-AUC for (c) eight classical classifiers and (d) five CNN models and logistic regression extended with nucleotide identities at positions +2 and +3, each trained using four alternative descriptors of amino acid substitutions while retaining all other predictors: Grantham distance (purple), FoldX ΔΔG (blue), sequence-based DDGun ΔΔG (red) and structure-based DDGun ΔΔG (green), with paired-bootstrap comparisons demonstrating equivalent performance within a margin of ±0.01 ROC-AUC for all comparisons except random forest, for which all three stability measures reduced performance relative to Grantham distance. (e) Sequence-context enrichment around TCA motifs, shown as the log2 ratio of nucleotide frequencies at mutated versus non-mutated sites.
